# The non-canonical SMC protein SmcHD1 antagonises TAD formation on the inactive X chromosome

**DOI:** 10.1101/342147

**Authors:** Michal R Gdula, Tatyana B Nesterova, Greta Pintacuda, Jonathan Godwin, Ye Zhan, Hakan Ozadam, Michael McClellan, Daniella Moralli, Felix Krueger, Catherine M Green, Wolf Reik, Skirmantas Kriaucionis, Edith Heard, Job Dekker, Neil Brockdorff

## Abstract

The inactive X chromosome (Xi) in female mammals adopts an atypical higher-order chromatin structure, manifested as a global loss of local topologically associated domains (TADs), and formation of two mega-domains. In this study we demonstrate that the non-canonical SMC family protein, SmcHD1, which is important for gene silencing on Xi, contributes to this unique chromosome architecture. Specifically, allelic mapping of the transcriptome and epigenome in SmcHD1 null cells revealed the appearance of sub-megabase domains defined by gene activation, CpG hypermethylation and depletion of Polycomb-mediated H3K27me3. These domains, which correlate with sites of SmcHD1 enrichment on Xi in wild-type cells, additionally adopt features of active X chromosome higher-order chromosome architecture, including partial restoration of TAD boundaries. Xi chromosome architecture changes also occurred in an acute SmcHD1 knockout model, but in this case, independent of Xi gene de-repression. We conclude that SmcHD1 is a key factor in antagonising TAD formation on Xi.

## Introduction

X chromosome inactivation is the mechanism that evolved in mammals to equalise levels of X-linked gene expression in XX females relative to XY males. Cells of early female embryos selectively inactivate a single X chromosome, usually at random, resulting in the formation of a stable heterochromatic structure, the Barr body. The inactive X chromosome (Xi), once established, is highly stable, and is maintained in somatic cells throughout the lifetime of the animal^1,2^. The X inactivation process is triggered by the non-coding RNA Xist, which localises to the Xi territory to induce chromosome-wide gene silencing^3–6^.

Chromatin features that distinguish Xi and the active X chromosome (Xa) include specific histone post-translational modifications, variant histones, and CpG DNA methylation (reviewed in ^2^). Additionally, Xi acquires a characteristic higher-order chromosome structure. Specifically, A-type chromatin compartments, corresponding to gene-rich regions which normally replicate in early S-phase, switch to replication in mid- or -late S-phase (reviewed in ^7^). Additionally, topologically associated domains (TADs), sub-megabase scale domains which are formed by the activity of cohesin, restricted at boundaries by oppositely oriented binding sites for the insulator protein CTCF^8–13^, are in large part absent on Xi, being replaced instead by two large mega-domains that are separated by a hinge that encompasses the DXZ4 repeat sequence^14–18^. The basis for this unique TAD structure is not well understood, but is thought to depend, at least in part, on ongoing expression of Xist RNA^17^.

Barr body formation is a multistep process. Thus, Xist RNA recruits specific chromatin modifiers, including the SPEN-NCoR-HDAC3 complex^19–22^, required for histone deacetylation^22^, and the PRC1 and PRC2 Polycomb complexes, required for deposition of H2A lysine 119 ubiquitylation (H2AK119u1) and H3 lysine 27 methylation (H3K27me3) respectively^23–27^. The lamin B receptor^22,28^, and m6A RNA modification complex^19,29^, have also been implicated in establishment of chromosome-wide gene silencing. Other factors are recruited to Xi at later stages. Examples include the variant histone macroH2A^30^, and the non-canonical SMC protein SmcHD1^31^. The role of these factors remains to be defined, although is likely to be linked to the long-term stability of the inactive state.

SmcHD1 is classified as an SMC protein by virtue of an SMC hinge domain at the C-terminal end, but differs from canonical SMC complexes in having a functional GHKL-ATPase domain rather that a Walker A/B type ATPase domain^32^. Biochemical and biophysical studies indicate that SmcHD1 homodimerises via the hinge and GHKL domains to form a complex that is reminiscent of bacterial SMC proteins, both in form and scale^33^, albeit forming a functional homodimer rather than a trimeric complex. SmcHD1 performs an important role in silencing on Xi, and at selected mono-allelically expressed autosomal loci ^31,32,34,35^. Whilst it is known that a proportion of Xi genes are activated in SmcHD1 mutant embryos^34,35^, the molecular mechanism is not well understood. Notably, although SmcHD1 is required for DNA methylation at CpG island (CGI) promoters of many Xi genes, loss of CGI methylation does not appear to account for the observed gene activation^34^. An alternative hypothesis is that SmcHD1-mediated compaction of Xi, inferred by microscopy based analyses in human cell lines^36^, imposes gene repression. Given the important role of SMC family proteins in genome topology, we set out to investigate the role of SmcHD1 in the higher-order architecture of Xi. Thus, we performed high-resolution analysis of Xi transcription, epigenetic features, and higher-order chromatin features in SmcHD1 null cell lines. Our findings demonstrate that SmcHD1 on Xi plays a role in transcriptional and epigenetic regulation of sub-megabase chromosome domains, and in antagonising TAD formation.

## Results

### SmcHD1 enriched sites on Xi show widespread Xi gene activation in SmcHD1 null MEFs

In order to gain insight into the role of SmcHD1 in X inactivation we set out to analyse epigenomic and long-range chromatin features of Xi at high resolution. Thus, we derived XX MEF lines from *Mus musculus domesticus* (domesticus) x *Mus musculus castaneus* (castaneus) female embryos, that were either wild-type (WT) or SmcHD1 null. Xi was of castaneus origin in both cases (Figure S1a,b). The high frequency of SNPs between domesticus and castaneus genomes allows assignment of high-throughput sequencing reads to either maternal or paternal genomes. The breeding strategy enabled us to obtain cell lines from F2 embryos in which the entire X chromosome was either of domesticus or castaneus origin. Autosomes on the other hand were mosaic as a result of recombination in the F1 generation. X inactivation in the interspecific embryos is random, so stable MEF lines were sub-cloned to obtain WT and SmcHD1 null lines. Xist RNA FISH and karyotype analysis confirmed the presence of Xi and Xa chromosome(s) (Figure S1c-e).

Initially, we performed allelic ChIP-seq analysis of SmcHD1 to define binding sites on Xi. As shown in Figure 1a, SmcHD1 is highly enriched over domains that correlate with gene rich regions across the length of the chromosome. We also observed SmcHD1 enrichment over specific regions on autosomes, including known SmcHD1 target loci such as the protocadherin locus on chromosome 18 and the PWS/AS imprinted gene cluster on chromosome 7 (Figure S2a,b). SmcHD1 peaks in the latter region accord with a previously published dataset analysing SmcHD1 occupancy in XY neuronal stem cells^37^.

**Figure 1.**
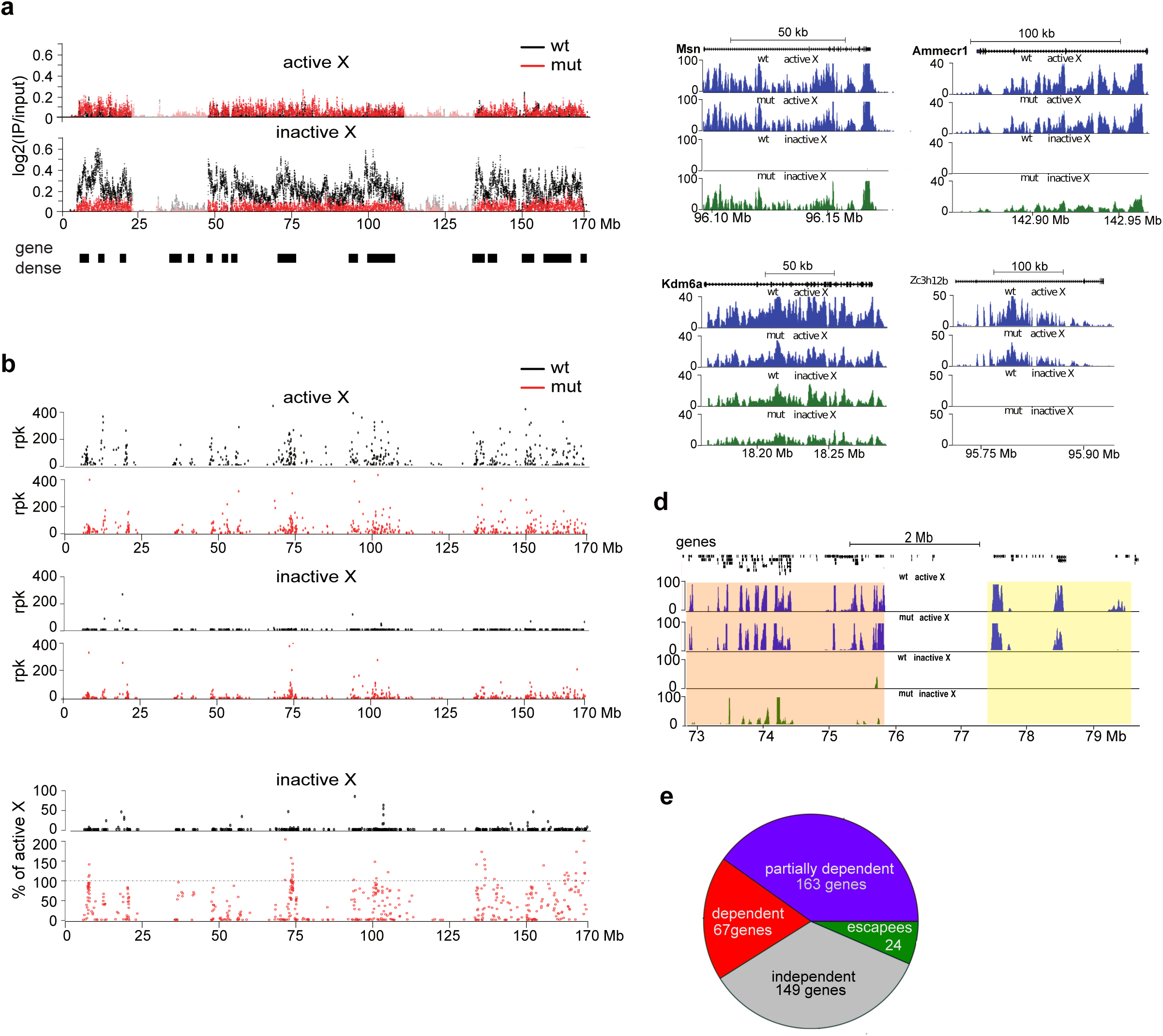
Widespread de-repression of Xi genes in SmcHD1 null MEFs. a. Chromosome-wide profiles of SmcHD1 occupancy on Xa and Xi depicted as allele-specific ChIP-seq log2 ratios of IP to input per 500 kb binned into 10 kb intervals. Profiles from WT and SmcHD1 null (mut) MEFs are presented. Black bars below the profiles depict gene-dense areas. b. Quantification of gene expression on Xa and Xi with allele-specific chromatin RNA-seq in WT and SmcHD1 null MEFs. Normalized read count per 1 kb of gene body is shown. Lower panel depicts gene expression on WT and SmcHD1 null Xi presented as % of Xa expression c. Representative examples of individual X-linked genes (UCSC screen shots) d. Example of a ∼ 7Mb region with a domain of strongly activated Xi genes, followed by a domain of Xi genes activated to a lesser degree (both with orange shading), and by a domain with genes expressed on Xa and silenced on Xi in SmcHD1 null cells (yellow shading). e. Summary of gene expression analysis. X-linked genes expressed on Xa were divided based on their transcriptional status on Xi into SmcHD1-dependent (expression not significantly different from Xa), partially SmcHD1-dependent (de-repressed but expressed significantly lower level than Xa), SmcHD1-independent (silenced on Xi) and WT escapees.

We went on to analyse Xi transcription in SmcHD1 null compared to WT cells using allelic chromatin RNAseq (ChrRNA-seq), which enriches for nascent unprocessed mRNAs, thereby maximising the number of informative SNPs due to inclusion of intron sequences. The results are summarised in Figure 1b-e. In WT cells, Xi expression was detected for a small number of genes (28/404), largely corresponding to genes previously reported to escape X inactivation (Figure S2c). However, in SmcHD1 null cells, Xi expression was seen for the majority of Xi genes (254/404) (Figure 1b-e). For 67 SmcHD1-dependent genes, Xi expression was in a similar range to that found on Xa, with a further 163 genes showing partial or low-level activation (Figure 1b-e, Figure S2c). We observed extensive overlap with 64 SmcHD1 dependent genes identified in a prior study in which we performed non-allelic microarray analysis of SmcHD1 null embryos^34^ (Figure S2d). The larger number of Xi expressed genes observed in this study can be attributed to the increased sensitivity afforded by allelic ChrRNA-seq.

Analysis of the association of SmcHD1-dependent genes with genomic sequence features in the immediate chromosomal environment identified a correlation with gene-rich regions, and the location of SINE repeats, as reported previously^34^. Because a relatively large proportion of Xi genes show SmcHD1 dependence, the latter correlation likely reflects the preferential location of SINE repeats within gene-rich regions. No other genomic features were found to correlate with SmcHD1 dependence.

SmcHD1 recruitment is a late step in Xist-mediated chromosome silencing^31^, suggesting it has a role in the continuation or maintenance of gene silencing. To further investigate this issue we generated an acute SmcHD1 null MEF line by CRISPR/Cas9-mediated mutagenesis in WT MEFs (Figure S3a-c). We then performed ChrRNA-seq as described above. Interestingly, and in contrast to embryonic SmcHD1 loss of function, no activation of Xi genes was observed (Figure S3d). This result suggests that SmcHD1 is required to reinforce gene silencing during a specific window in development, and is then dispensable, presumably reflecting compensation through other maintenance pathways.

### Unique features of the Xi epigenome in SmcHD1 null MEFs

Previous work established that SmcHD1 is important for DNA methylation of Xi CGI promoters^31^, but its role in chromosome-wide intergenic and intragenic DNA methylation on Xi has not been investigated. To address this we used whole genome bisulfite analysis (WGBS) to determine the methylome of Xa and Xi at single nucleotide resolution in WT and SmcHD1 null MEFs. A summary of data illustrating overall methylation density in 100kb windows across the entire genome is shown in Figure 2a. CpG methylation across autosomes was generally in the range of 60-80%, but was significantly lower on the X chromosome. We noted that gene-poor regions are CpG hypomethylated across all chromosomes. An example, chromosome 7 is illustrated in Figure S4a. A similar pattern of hypomethylation is apparent in available MEF WGBS^38^ (Figure S4b). Accordingly, we find that total CpG methylation levels in MEFs, as determined by HPLC analysis, are moderately reduced relative to ES cells and adult tissue (Figure S4c). The molecular basis for this pattern of CpG hypomethylation is currently unknown.

**Figure 2.**
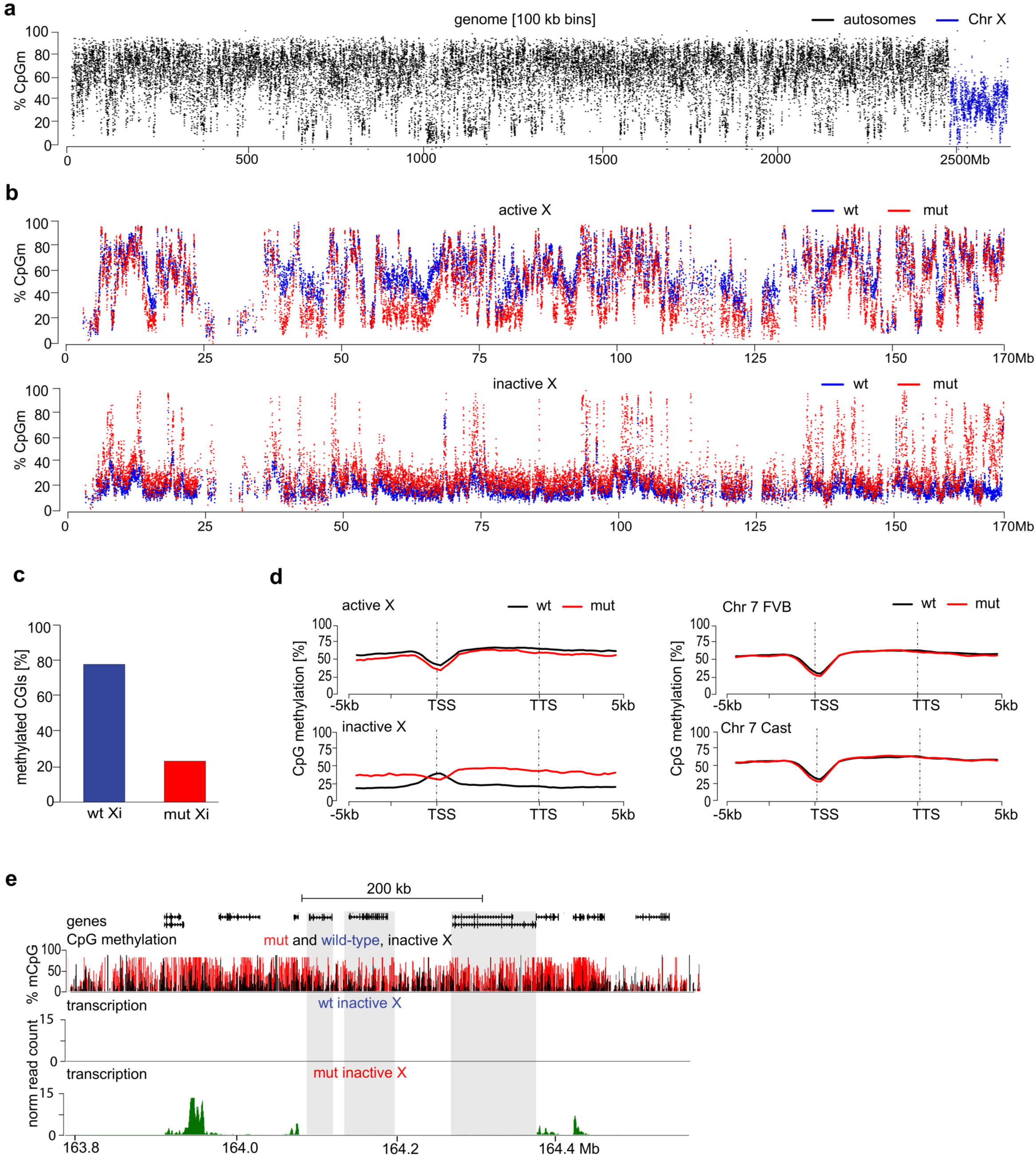
The Xi methylome in WT and SmcHD1 null MEFs. a. WGBS reads binned into 100 kb intervals illustrating reduced level of CpG methylation on the X chromosome relative to autosomes. b. Allele-specific DNA methylation profile of Xa (top) and Xi (bottom) in WT and SmcHD1 null MEFs plotted as % of CpGm averaged within 10 kb bins. c. Fraction of CGIs methylated on Xi in WT and SmcHD1 null MEFs. A total of 119 CGIs were analyzed for which at least 5 cytosines were covered by at least 3 reads in all allele-specific alignments. CGIs with mean methylation level >=40% were classified as methylated <40% as unmethylated. d. Metagene plots of DNA methylation in WT and SmcHD1 null MEFs generated for Xa and Xi as well for maternal and paternal Chr 7. e. Example of a hyper-methylated domain on Xi in SmcHD1 null cells, showing that the region of high DNA methylation extends beyond expressed genes to encompass genes that are silenced. Non-expressed genes with DNA methylation above the background level are highlighted with shadowed boxes.

We went on to compare allelic methylation levels on Xa and Xi chromosomes (Figure 2b). In WT MEFs, Xa methylation patterns were similar to autosomes (56% compared with 66% CpG methylation), but Xi was extensively CpG hypomethylated across the entire chromosome (19% CpG methylation). CpG hypomethylation of gene-poor regions, as described above, likely contributes to the observed pattern on Xi. However, CpG hypomethylation was also evident in gene-rich regions, presumably linked to X inactivation status. This observation is consistent with prior studies showing reduced levels of CpG hypomethylation on Xi relative to Xa^39,40^. Xi CGIs in WT MEFs were, as expected, highly methylated (Figure 2c). Additionally, we observed high levels of CpG methylation over bodies of genes that escape X inactivation (Figure S4d).

In SmcHD1 null MEFs, Xi CGIs were in most cases hypomethylated (Figure 2c), as previously reported^31^. Conversely, we observed several domains with relatively high CpG methylation (Figure 2b). These domains correspond in most cases with Xi genes that are activated in SmcHD1 null MEFs, although CpG hypermethylation is not restricted to transcribed sequences, extending both 5’ and 3’ (Figure 2d,e). Individual genes that normally escape X inactivation were also CpG hypermethylated on Xi, similar to WT MEFs (Figure S4d).

The histone modification H3K27me3 catalysed by the major Polycomb complex PRC2, is highly enriched on Xi as determined by immunostaining^23,24^, and high-resolution ChIP-seq analysis^41^. Previously we noted that this feature is not grossly affected by SmcHD1 loss of function, as determined by immunostaining of interphase nuclei in XX embryos^32^. Similarly, H3K27me3 enrichment on Xi was readily detected by immunostaining in the WT and SmcHD1 null MEF lines described herein (Figure S3b). However, allelic ChIP-seq analysis revealed domains in which Xi H3K27me3 is markedly depleted in SmcHD1 null cells (Figure 3a). These domains (estimated 77 with mean length of 140Kb), comprise approximately 10% of the Xi regions over which H3K27me3 is normally enriched. The location of the H3K27me3 depleted domains correlates closely with CpG hypermethylation and Xi gene activation (Figure 3b,c). However, as noted for CpG hypermethylation domains, the H3K27me3 depleted regions extend beyond the boundaries of activated genes (Figure S5a), and in some cases do not include known genes (Figure S5b). Together these results suggest that modified epigenomic features in SmcHD1 null cells correlate with domains in which Xi genes are activated.

**Figure 3.**
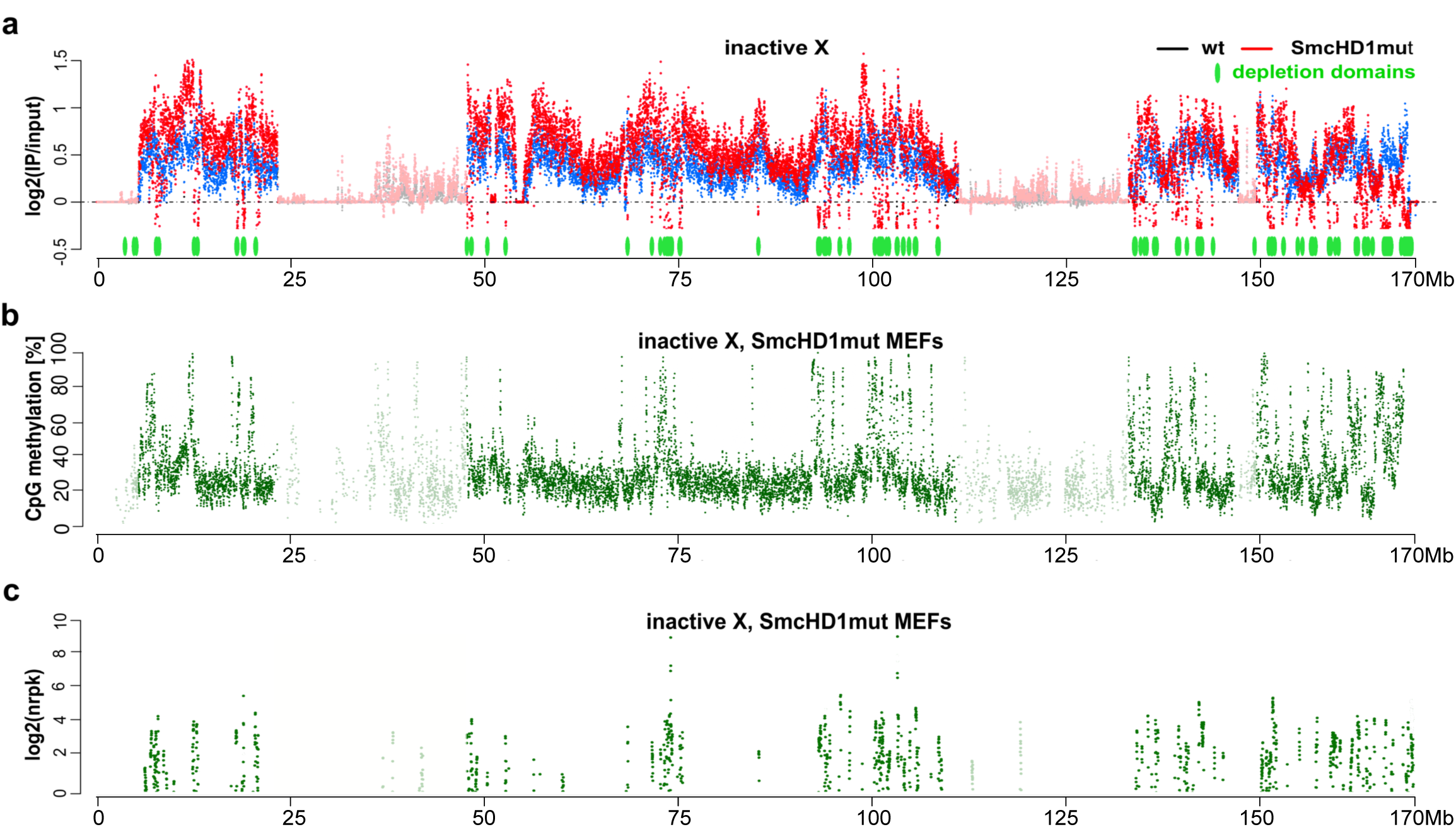
H3K27me3 depletion highlights domains of Xi gene activation in SmcHD1 null MEFs. a. Allele-specific H3K27me3 ChIP-seq illustrating multiple H3K27me3 depleted domains not present on WT Xi. Scatterplot depicts ChIP log2(IP/input) values averaged within 10 kb bins. 77 H3K27me3 depleted domains are plotted in green below the H3K27me3 profile. Regions of low probe mappability are indicated with reduced transparency. Location of H3K27me3 depleted domains on Xi in SmcHD1 null MEFs correlates with, CpG hypermethylation domains (b) and genes activated on Xi (c).

### A modified higher-order chromosome architecture on Xi in SmcHD1 null cells

In light of evidence that SmcHD1 influences Xi chromatin at the level of sub-megabase domains, we went on to directly analyse parameters of long-range chromatin architecture. Mammalian chromosomes are comprised of distinct gene-rich and gene-poor compartments with sizes ranging from sub-megabase to several megabases in length. These regions are replicated co-ordinately, either during early- or mid/late-S phase respectively. Replication timing domains are broadly synonymous with A- and B- type chromatin, which in studies on long-range chromosome topology, have been shown to self-associate^42^. Additionally, B-type domains overlap extensively with Lamin associated domains (LADs) which localise to the nuclear periphery^43^. Xi is unusual in that both gene-rich and gene-poor compartments replicate synchronously in mid/late S-phase^7^.

A previous analysis of a human XX somatic cell line in which SMCHD1 was depleted using siRNA, revealed an aberrant replication timing pattern for Xi, with the appearance of early replicating domains as determined using a cytogenetic assay^36^. With this observation in mind we set out to determine allelic temporal replication patterns in our WT and SmcHD1 null MEFs using RepliSeq, a high-resolution sequencing based approach^44^ (Figure S6a,b). In WT MEFs we observed mid/late S-phase replication across Xi, contrasting with Xa where gene-rich and gene-poor chromosome domains replicate in early- and mid/late S-phase respectively (Figure 4a). However, in SmcHD1 null MEFs, replication patterns on Xi are more similar to the Xa pattern in WT cells (Figure 4b). Thus, we observed regions of the chromosome in which Xi replication timing overlaps with that seen on Xa (orange shading, Figure 4b), and other regions where replication timing is either partially advanced or largely unaffected (yellow shading, Figure 4b). The location of Xi regions showing a more pronounced shift in replication timing correlates closely with domains of H3K27me3 depletion (Figure 4c), a proxy for SmcHD1 localisation, Xi gene activation and CpG hypermethylation, as above.

**Figure 4.**
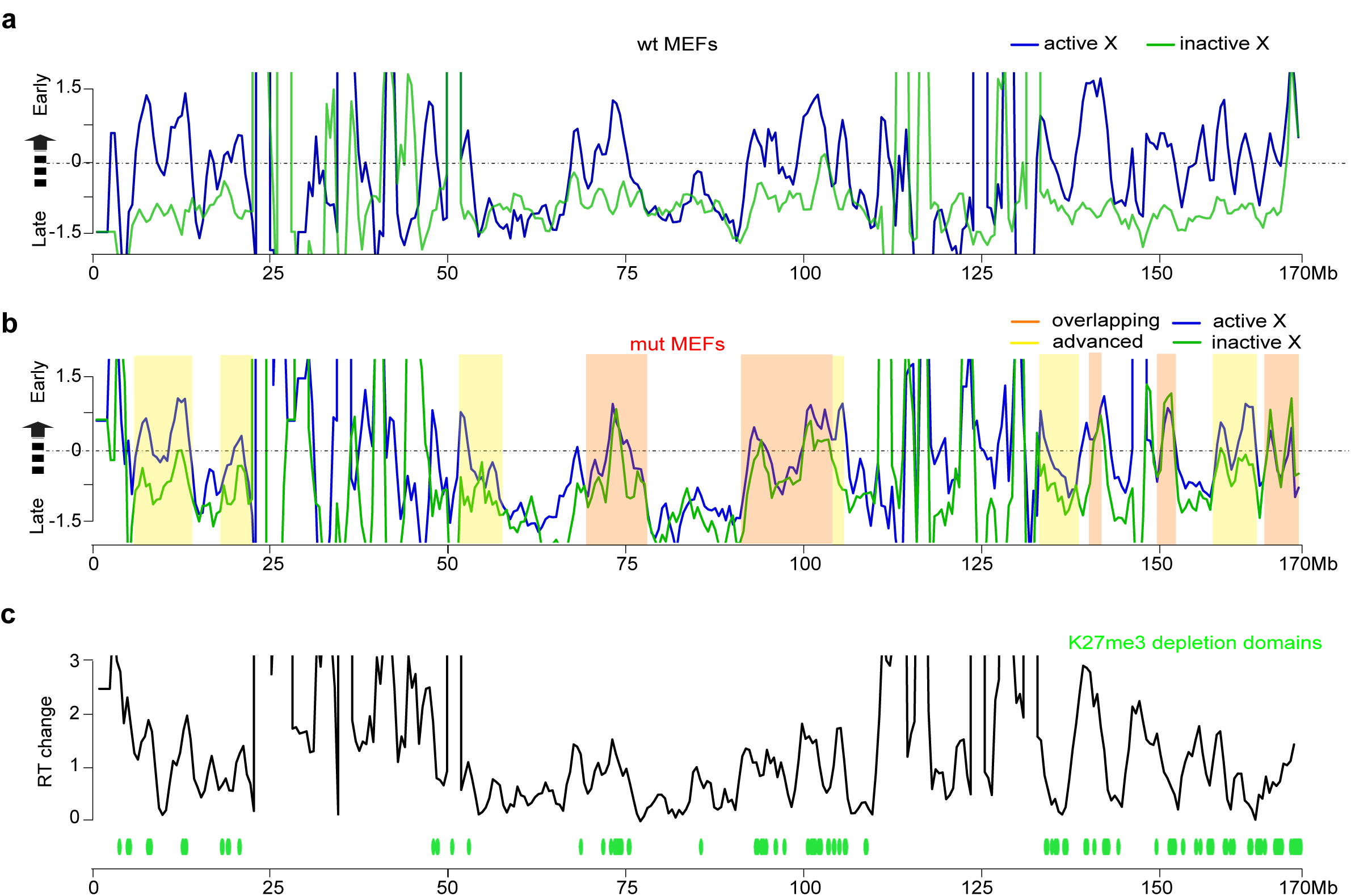
Advanced Xi replication timing in SmcHD1 null MEFs. Allele-specific Repli-seq profiles of WT (a) and SmcHD1 null (b) MEFs. Z-score of ratio of read densities from asynchronously dividing cells in S-phase to the read densities of G1 cells, 500 kb bins. Higher Z-score value corresponds to earlier replication. Shaded areas correspond to regions with a high density of unmappable reads. Boxes represent examples of regions in which replication timing on Xi and Xa in SmcHD1 null cells are equivalent (orange), or moderately advanced/equivalent to WT Xi (yellow). Regions of low probe mappability are indicated with reduced transparency. c. Difference in replication timing (Z-score of G1/S read densities) of Xa and Xi in WT and SmcHD1 null MEFs. Xi H3K27me3 depletion domains are indicated in green at the bottom of the plot.

A further level of chromosome architecture is TADs, to which replication timing domains are thought to be linked^45^. To determine if TAD organisation on Xi is affected by SmcHD1 loss of function we performed Hi-C in the SmcHD1 null and WT MEF cell lines. Analysis of data for autosomes and for Xa indicated that SmcHD1 does not impact on global TAD structure. An example, chromosome 7, is illustrated in Figure S7a, b. Along the Xi in WT cells we detect the presence of two megadomains separated by the DXZ4-containing boundary and the general absence of TADs, as previously reported^14–18^ (Figure 5). However, we observed several distinct features in SmcHD1 null cells. The mega-domain structure and DXZ4 hinge characteristic of Xi are discernible, but we observed a reduction in long-range interactions (Figure 5a-c). Additionally, at several sites we observed restoration of Xa TAD structure (Figure 5d, Figure S7c). In the most distal 20Mb of Xi and Xa TADs and their boundaries are virtually indistinguishable (Figure 5d). TAD restoration correlates with the sites of Xi gene activation and modified CpG methylation/H3K27me3 patterns (Figure 5e, Figure S7d). Analysis of interaction frequencies further illustrates the increase in short-range interactions in SmcHD1 null MEFs, with variation across the chromosome correlating with transcriptional and epigenome changes (Figure 5f, Figure S7e). Taken together these results demonstrate that SmcHD1 plays an important role in defining the unique higher-order chromatin domain organisation on Xi.

**Figure 5.**
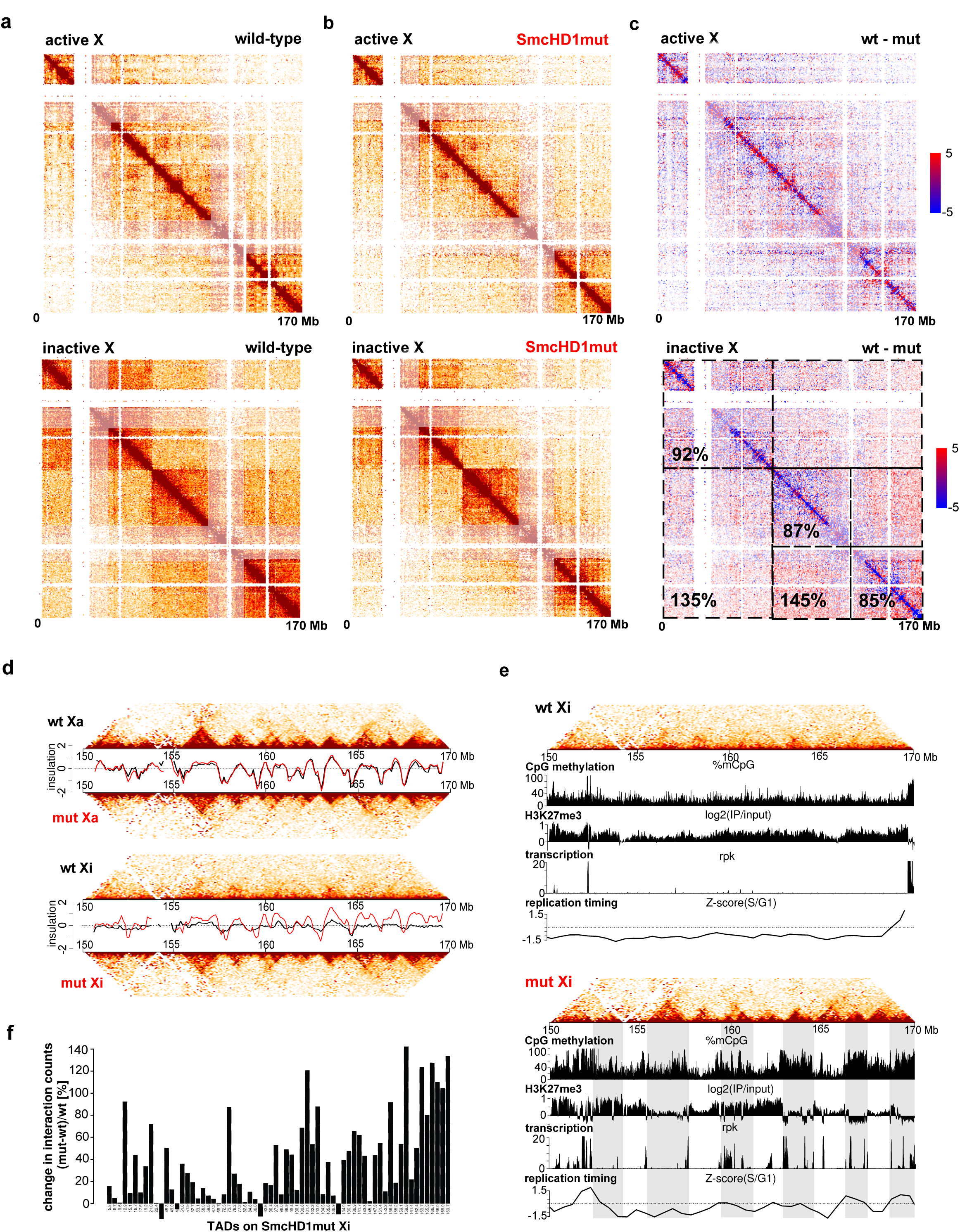
Altered TAD structure on Xi in SmcHD1 null MEFs. a. Heatmap depicting allele-specific Hi-C interactions for Xa and Xi for WT MEFs. Interaction matrices were generated from Hi-C interactions normalised for the number of available reads per chromosome, ICE-balanced and binned into 500 kb bins. b. Heatmap with Hi-C interactions for Xa and Xi in SmcHD1 null MEFs, data processed as above. c. Difference between normalised interaction counts of WT and SmcHD1 null MEFs. Heatmap for Xi presents additionally quantification of differences in interaction counts between WT and SmcHD1 null MEFs for distinct parts of the Chr X interaction matrix d. Local chromatin conformation of the distal 20 Mb of Xa and Xi for WT and SmcHD1 null MEFs. Heatmaps were generated as above, but interactions were binned into 100 kb bins. Interaction directionality was quantified by isolation score plotted in black for WT and in red for SmcHD1 null MEFs. e. Allele-specific Hi-C heatmaps of Xi in WT (top) and SmcHD1 null (bottom) MEFs, aligned with DNA methylation, H3K27me3, transcription and replication timing profiles for the distal 20 Mb of Chr X. Shaded bars highlight TAD borders for SmcHD1 null cells. f. Comparison of the interaction counts from distinct TADs on the Xi of SmcHD1 null MEFs with the interaction counts within the same intervals on the WT Xi. The values were calculated as the difference between the counts in mutant and WT MEFs related to the count number in WT MEFs and presented as %. Interactions over 100 kb falling within TAD borders were scored.

### SmcHD1 depletes CTCF and cohesin at TAD boundaries on Xi

Recent studies have established that TAD boundaries result from the insulator protein CTCF restraining processive activity of the cohesin complex^11–13^. Accordingly, CTCF and cohesin occupancy is reduced at many sites on Xi compared with Xa^17,46,47^. With this in mind we performed ChIP-seq to determine the occupancy of CTCF and the cohesin subunit, Rad21, on Xa and Xi in WT compared with SmcHD1 null MEFs. Consistent with these prior studies, we observed that both CTCF and Rad21 levels are depleted on Xi compared with Xa (Figure 6a-e). In contrast, in SmcHD1 null MEFs we observed restoration of both CTCF and Rad21 occupancy at many sites (Figure 6a-c). This effect is most apparent over chromosomal regions at which SmcHD1 is enriched in WT cells, and at which epigenome and chromosome architecture changes occur in SmcHD1 null cells (Figure 6a). At selected regions for which allelic assignment of CTCF/Rad21 enrichment sites was possible, we were able to correlate restoration of CTCF/Rad21 occupancy on Xi with the re-appearance of specific TADs (Figure 6c). The majority of Xa CTCF/Rad21 binding sites that were absent on Xi in WT MEFs acquired significant levels of CTCF/cohesin occupancy in SmcHD1 null cells (Figure 6d,e).

**Figure 6.**
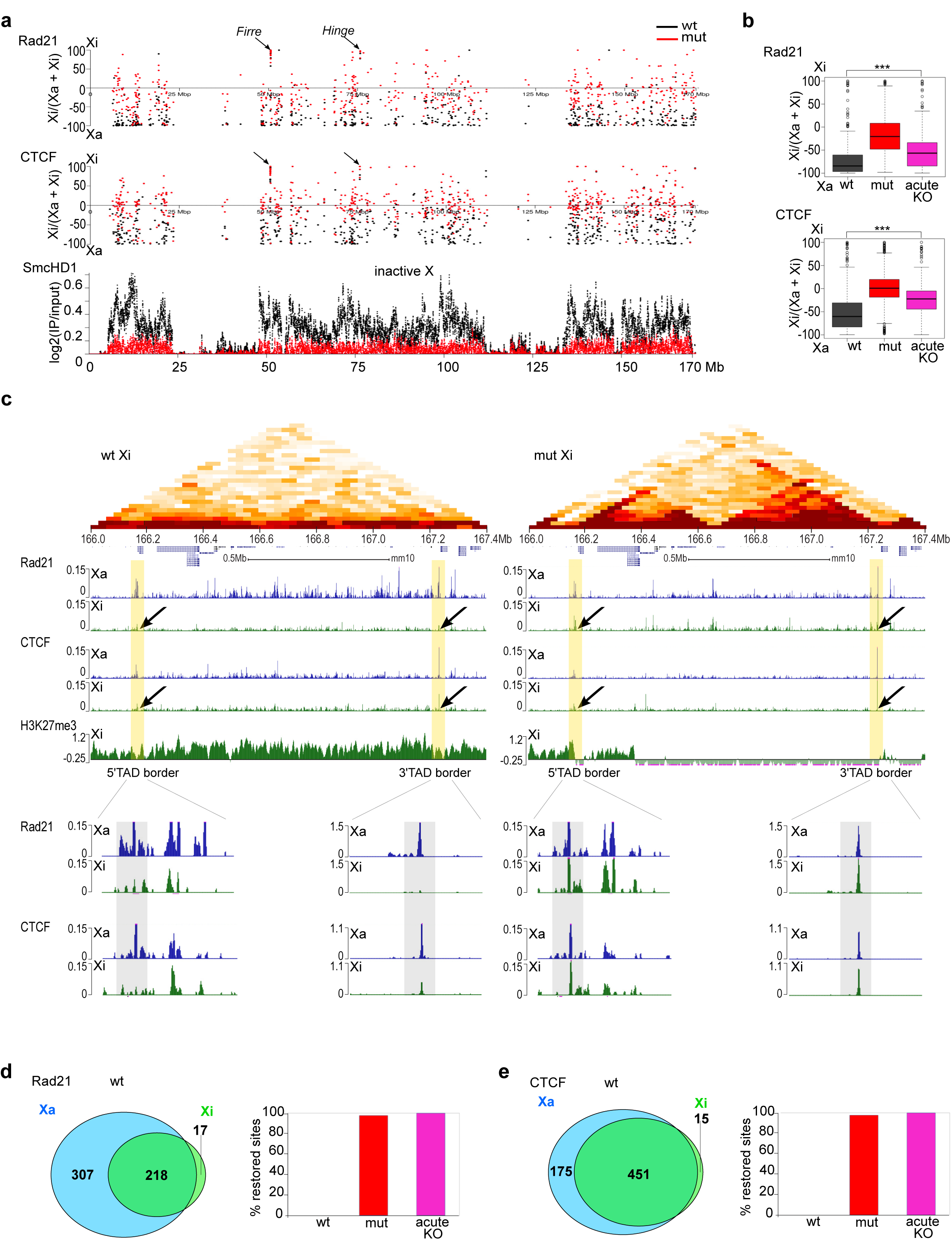
CTCF and cohesin occupancy on Xi in SmcHD1 null cells. a. X Chr-wide occupancy of Rad21 (top panel) and CTCF (middle panel) for Xi or Xa, estimated for distinct peaks occupied WT, SmcHD1 null and acute SmcHD1 null cell lines, and shown as a ratio indicating relative levels on Xa and Xi. 100=fully Xi specific, −100=fully Xa specific. Black arrows depict Firre and Hinge loci which have high occupancy on WT Xi. Lower panel, for reference, shows SmcHD1 profile on the Xi log2(IP/input) in 10 kb bins. b. Summary of preferential occupancy of Rad21 (top panel) and CTCF (bottom panel) for Xi or Xa, estimated for the peaks occupied in all three cell lines (WT, SmcHD1 null and acute SmcHD1 null). Boxplots present quartiles, median and outliers of re-scaled ratio of Xi specific peaks/(sum of the peaks from both chromosomes). c. Example illustrating Hi-C interactions, CTCF and Rad21 occupancy within a region harbouring a TAD that is re-established in SmcHD1 null cells. Top panel shows HiC heatmap of chromatin interaction counts in 50 kb bins. Black arrows depict occupancy changes within the borders of the re-established TAD. Yellow shading highlights regions which differ in CTCF/Rad21 binding between KO and WT cells which correlate with external borders of the re-established TAD.Lower panels show enlarged views of 5’ and 3’ TAD borders. d,e. Venn diagram quantifying Rad21 (d) and CTCF (e) occupancy on Xi and Xa for peaks occupied in WT, SmcHD1 null and acute SmcHD1 null cell lines. Peaks were considered Xi- or Xa- specific if >90% of all reads overlapping a distinct peak map to Xi or Xa respectively. The remainder of peaks were counted as being present on both alleles. Barplot shows the proportion of the Xa-specific peaks which gained Rad21 (d) and CTCF (e) in SmcHD1 null (red) and acute SmcHD1 null (magenta) MEFs.

To assess whether changes in CTCF/Rad21 occupancy depend on Xi gene activation, we analysed the acute SmcHD1 null MEF line in which there is no detectable Xi gene de-repression. Interestingly, restoration of CTCF/Rad21 occupancy on Xi was also evident in this cell line (Figure S8a,b), although Rad21 occupancy was of a lesser magnitude compared to MEFs derived from SmcHD1 null embryos (Figure 6b,d,e). This result suggests SmcHD1 directly affects TAD formation on Xi, independent of its role in maintaining Xi gene silencing.

## Discussion

Allelic ChIP-seq analysis of SmcHD1 on Xi shows strong enrichment over gene-rich domains. This pattern mirrors the localisation of Xist RNA and Xist-dependent histone modifications^48,49^, and is therefore consistent with previous studies that reported a requirement for ongoing Xist expression to maintain SmcHD1 enrichment on Xi^36^. Using allelic RNA-seq we found that a large proportion of Xi genes within SmcHD1 enriched regions are de-repressed in SmcHD1 null MEFs, albeit, in most cases expression does not reach the level seen on Xa. At present we cannot discriminate between the possibility that Xi gene activation in SmcHD1 null cells occurs in a graded fashion within individual cells, or that low level activation reflects probabilistic expression in a subset of cells that varies for different loci.

In prior work we found that SmcHD1 enrichment on Xi is a late step in the X inactivation cascade in differentiating XX ESCs^31^, suggesting a role in maintenance rather than establishment of X inactivation. Paradoxically, in this study we found that there is no activation of Xi genes following acute knockout of SmcHD1 in MEFs. In light of this we speculate that SmcHD1 is required for Xi to transition from the initial repressed state to longterm silencing, but that downstream silencing pathways, for example CGI DNA methylation, maintain the inactive state even in the absence of SmcHD1. Although we cannot at present define how SmcHD1 mediates transitional Xi gene silencing, its role in establishing higher-order chromosome folding, discussed below, is one possible mechanism.

Analysis of the Xi methylome at single nucleotide resolution confirms prior studies demonstrating Xi hypomethylation^39,40^. We observed widespread and extensive hypomethylation, encompassing both gene-rich and gene-poor compartments. As noted, we cannot rule out that hypomethylation of gene-poor regions on Xi is linked to genome-wide effects in MEFs. Hypomethylation of gene-rich regions on Xi on the other hand, is presumably linked to X inactivation status. Mechanistically, this could be attributable to reduced binding of the de novo DNA methyltransferases Dnmt3a/b, which recognise the histone modification H3K36me3 in gene bodies of active genes^50,51^. Consistent with this idea, hypermethylated regions on Xi correspond to genes that escape X inactivation and in SmcHD1 null MEFs, hypermethylation occurs within regions encompassing activated Xi genes. However, Xi CpG hypermethylation in SmcHD1 null MEFs is not restricted to gene bodies, where H3K36me3 is normally enriched, but rather occurs across domains that include upstream and downstream sequences, and in some cases more than one gene. Reciprocal depletion of PRC2-mediated H3K27me3 further reinforces that deregulation of the Xi epigenome in SmcHD1 null cells occurs at the level of sub-megabase domains. These observations suggest alternative mechanisms, for example increased accessibility to nucleosome remodelling complexes, are involved in establishing the distinct epigenomic features observed on Xi in SmcHD1 null MEFs.

Further support for SmcHD1 functioning on Xi at the level of domain organisation comes from direct analysis of replication-timing domains and TADs. In both cases we observe a shift towards Xa-specific organisation across Xi, most obviously associated with sites where Xi gene activation and modified epigenomic features occur. The mechanism for replication-timing changes is not known. One possibility is that SmcHD1 affects recruitment of Rif1, required to direct PP1 to reverse Cdc7-mediated phosphorylation of the MCM complex^52^. Thus, at chromosomal regions that are normally early replicating, SmcHD1 may facilitate Rif1 recruitment, for example through affecting chromosome architecture. SmcHD1 has been reported to disrupt Xi replication-timing independent of gene activation following SMCHD1 knockdown in a human XX cell line^36^, implying that transcriptional activation is not the cause of shifted replication-timing patterns.

The unique organisation of TADs on Xi has been linked to reduced binding of CTCF at TAD boundaries and/or reduced recruitment of cohesin complexes^17,18^. Accordingly, we observe restoration of both CTCF binding and cohesin (Rad21) accumulation in SmcHD1 null MEFs, and sites of restoration correlate well with the reappearance of Xa TAD structure. These findings accord with prior analysis of SmcHD1 at an autosomal target, the protocadherin gene cluster in neural stem cells, which suggested that SmcHD1 and CTCF have opposing roles in transcriptional regulation^37^. The antagonistic effects of SmcHD1 on TAD boundaries may be analogous to the role of condensin in TAD dissolution during mitosis^53^.

We observed that whilst restoration of CTCF binding also occurs in MEFs following acute deletion of SmcHD1 (in which there is no Xi gene de-repression), restoration of Rad21 on Xi was of a lower magnitude. Prior work has suggested a link between cohesin loading and active transcription^54^, and this may account for the reduced level of restoration relative to the chronic SmcHD1 null model. Regardless, this result indicates that changes in the long-range architecture of Xi in SmcHD1 null cells is not a consequence of gene de-repression. We note that prior studies have reported that depletion of Xist RNA in somatic cells leads to partial restoration of Xa chromosome architecture/TAD structure in the absence of changes in Xi gene repression^17,55^. Given that SmcHD1 recruitment to Xi is dependent on ongoing Xist expression^36^, we suggest that these effects may be attributable in part, or entirely, to loss of SmcHD1 on Xi.

In summary, our findings illustrate that SmcHD1 functions in X inactivation at the level of chromatin domain organisation. In future studies it will be important to determine the molecular interactions that underpin SmcHD1 function in antagonising TAD formation both on Xi and potentially at other SmcHD1 target loci.

## Acknowledgements

We thank members of the Brockdorff and Klose labs for helpful discussions and advice. We would like to thank Vadimir Benes and EMBL Gene Core for ChrRNA-seq and WGBS, Amanda Williams from Oxford University Zoology Dept. sequencing, the High-Throughput Genomics Group at the Wellcome Trust Centre for Human Genetics (funded by Wellcome Trust grant reference 090532/Z/09/Z and MRC Hub grant G0900747 91070) for the generation of the H3K27me3 Sequencing data, Miguel Branco for advice on whole genome bisulfite sequencing, and Michal Maj, Dunn School Flow Cytometry Facility for assistance in flow-sorting for Repli-seq. N.B. is funded by Wellcome (103768) and the European Research Council (340081). J.D. is an investigator of the Howard Hughes Medical Institute and was funded in part by a grant from the National Human Genome Research Institute (HG003143). SK is funded by Ludwig Cancer Research and BBSRC grant BB/M001873/1.

## Author contributions

N.B, J.D., E.H., M.R.G. and T.B.N. conceived this study. T.B.N. and M.R.G. performed most of the experimental work and data analysis. G.P., Y.Z. and H.O. contributed to Hi-C analysis. J.G. contributed to mouse genetics and cell line derivation. S.K., W.R. F.K. and M.M. contributed to DNA methylation analysis, C.M.G and D.M. performed karyotype analysis. N.B., T.B.N. and M.R.G. prepared the manuscript with input from all other authors.

## Methods

### Derivation and culture of mouse embryonic fibroblasts (MEFs)

Animal studies were carried out under the United Kingdom Home Office ASPA project, licence numbers 30/2800 (until 2015) and 30/3326 (from 2015 to present).

Interspecific crosses between SmcHD1 mutation carrier strain (FVB) and castaneus strain were designed to prevent meiotic recombination between X^cast^ and X^FVB^ chromosomes (Fig.1). FVB MommeD1^32^ heterozygous male was crossed with castaneus WT female. F1 SmcHD1 heterozygote male was selected for F2 cross with SmcHD1 heterozygous FVB female. Embryos from this cross were dissected at E9.5, and MEF lines were derived by culturing trypsinised WT and SmcHD1^mut/mut^ littermate embryos. Established MEF lines were subcloned, and individual clones were selected for further analysis. SmcHD1 genotype was determined by sequencing PCR fragments overlapping the SmcHD1 point mutation in exon 23^32^. FVB or castaneus origin of the X chromosome was determined by sequencing PCR fragments across Xist SNP region. XX/XY genetic content was established by PCR analysis using primers that give distinct bands for the Uba1 and Uba1y genes^56^. All primers used for genotyping are listed below.

Fibroblasts were grown in Dulbecco’s modified Eagle medium (DMEM, Life Technologies) supplemented with 10% fetal calf serum (FCS; Seralab), 2mM L-glutamine, 1x nonessential amino acids, 50µM 2-mercaptoethanol, and 100U/ml penicillin/100µg/ml streptomycin (Life Technologies) in a humidified 37°C incubator under 5% CO2.

### CRISPR/Cas9-mediated SmcHD1 knockout in MEFs

sgRNAs targeting SmcHD1 in the vicinity of the original MommeD1 mutation (exon 23) and neighbouring exons were cloned into plasmid pX459 as described^57^. SmcHD1 wt1A2 cell line that carries WT SmcHD1 (two FVB alleles and one castaneus allele) was used for mutagenesis. Fibroblasts were plated on 90mm Petri dishes a day before transfection. A few hours before transfection growth medium was replaced with antibiotic-free medium. Cells were transfected with a pool of two sgRNAs, 3µg each, using Lipofectamine 3000 (Life Technologies) according to the manufacturer’s instructions. DNA:Lipofectamine ratio was 1:3. Cells were trypsinised 18hrs after lipofection and plated on 145mm petri dishes at densities 1/10; 1/3 and the rest. Puromycin selection at final concentration 4μg/ml was applied next day and maintained for 72hrs, after which cells were grown in EC10 medium for 10 days until colonies were ready to be picked. Individual well-spaced colonies were scraped from the dishes and transferred into 48-well plate without trypsinisation. Initial screening of knockout clones was by genomic PCR across intronic region (SmcHD1_TNK208+SmcHD1_TNK291). Selected clones were subsequently analysed by sequencing of cloned PCR products from regions around sgRNAs and across introns when appropriate. Candidate KO clones D4 and A3.3 were subsequently further characterised by immunofluorescence with SmcHD1, H3K27me3 and H2Aub1 antibodies, and by Western blot of nuclear extracts with SmcHD1 antibody.

### Metaphase spreads

Cells were plated on 140mm Petri dishes two days before metaphase collection. Semi-confluent cultures with actively dividing cells were fed with fresh medium supplemented with 1.5μg/mL ethidium bromide (Life Technologies) and incubated for 1hr 20min. Mitotic cells were arrested by addition of KaryoMAX Colcemid (Life Technologies) at a final concentration of 0.1μg/ml for further 40min. Cells were carefully rinsed once with PBS and trypsinised briefly at room temperature until the top layer of cells enriched with mitotic cells started to move. Trypsin was inactivated with fresh medium and top layer of cells was collected and pelleted at 200g for 3min, RT. All supernatant was aspirated and pellets were gently resuspended in 1ml of hypotonic solution (75mM KCl). After incubation for 5 min at RT, swollen cells were placed on ice, and 200μL of freshly prepared methanol/acetic acid fixative (3:1, 4°C) was added drop-wise to the solution to pre-fix cells. Cells were pelleted at 200g for 3min, and most of the supernatant was removed, leaving behind about 100μL to re-suspend the cells in by gentle flicking. 1ml of ice-cold fixative was added to the resuspended cells and incubated overnight at 4°C without agitation. The following day the cells were carefully resuspended in the same fixative and pelleted as before. The fixative was replaced 3-5 times in total until good metaphase chromosome spreading was observed after the cell suspension was dropped onto microscope slides and air dried.

### M-FISH

Metaphase spreads were prepared as described above, omitting the addition of ethidium bromide. The cells were hybridised with the 21XMouse MFISH kit (Zeiss Metasystem), following the manufacturer instructions. The slides were mounted in DAPI/Vectashield mounting medium (Oncor), under a glass coverslip, and analysed with an Olympus BX60 microscope for epifluorescence equipped with a Sensys CCD camera (Photometrics, USA). Images were collected using Genus Cytovision software (Leica). A minimum of twenty-five cells were analysed for each cell line.

### Chromosome paint (DNA FISH) on metaphase spreads

Slides with freshly fixed metaphase spreads were dehydrated through ethanol series (2×70%, 2×90% 2 min each followed by 100% for 5 min), and incubated in an oven at 65°C for 1hr. Slides were cooled down on a bench for a few minutes and denatured in 70% (v/v) Formamide/2xSSC at 65°C for 90sec. Slides were quenched in ice-cold 70% ethanol for 4 min and then dehydraded again through the ethanol series (the same as above), and dried in the vacuum dessicator for 5 min at RT. A 1:1 mix of directly labelled chromosome 8 (Cy-3, Cambio Ltd) and X (FITC) paints was denatured at 65°C for 10min, spun down and incubated at 37°C for 40 to 60min. Probe was incubated with the denatured metaphases overnight at 37°C. Next day the slides were washed twice with a solution of 1xSSC/50% formamide followed by two washes with 1xSSC and three washes with 4xSSC/0.05% Tween-20 in a water bath at 45°C, 5 min each. Slides were mounted in Vectashield containing 4,6-diamidino-2-phenylindole (DAPI) (Vector laboratories) and sealed with nail varnish.

### RNA-FISH

Cells were plated on Superfrost Plus gelatinised slides (VWR) and grown at least overnight. After washing twice with PBS, cells were permeabilised for 5 min in CSK buffer (100mM NaCl, 300mM sucrose, 3mM MgCl2, 10mM PIPES) with 0.5% Triton X-100 (Sigma) on ice. Slides were rinsed briefly in PBS and fixed in 4% formaldehyde/PBS for 10 mins on ice, followed by two washes in 70% ethanol. Slides were either stored in 70% ethanol at 4°C until use or dehydrated (80%, 95%, 100% ethanol, 3 min each, RT) and air dried immediately before hybridisation with Xist probe. Xist probe was generated from an 18 kb cloned cDNA spanning the whole Xist transcript using a nick translation kit (Abbott Molecular) as previously described^19^. Directly labelled probe (1.5μL) was co-precipitated with 10μg salmon sperm DNA, 1/10 volume 3M sodium acetate (pH 5.2) and 3 volumes of 100% ethanol. After washing in 75% ethanol, the pellet was dried, resuspended in 6μL deionised formamide and denatured at 75°C for 7 min before quenching on ice. Probe was diluted in 6μL 2x hybridisation buffer (5x SSC, 12.5% dextran sulfate, 2.5mg/mL BSA (NEB)), added to the denatured slides and incubated overnight at 37°C in a humid chamber. After incubation, slides were washed three times with a solution of 2xSSC/50% formamide followed by three washes with 2xSSC in a water bath at 42 °C. Slides were mounted with Vectashield with DAPI and sealed with nail varnish.

### Immunofluorescence

Cells were plated on Superfrost Plus gelatinised slides (VWR) at least a day before the experiment. On the day of the experiment, cells were washed with PBS and then fixed with 2% formaldehyde in PBS for 15 mins at RT, followed by 5 mins of permeabilisation in 0.4% Triton X-100. Cells were rinsed with PBS three times, 2 min each and pre-blocked with a 0.2% w/v PBS-based solution of fish gelatine (Sigma) 3 times, 5 min each. Primary antibody dilutions were prepared in fish gelatin solution with 5% normal goat serum. Primary antibody dilutions are listed below. Cells were incubated with primary antibodies for 2 hours in a humid chamber at room temperature, then washed three times in fish gelatin solution to remove non-bound and non-specifically bound antibodies. Secondary antibodies were diluted in fish gelatin solution and incubated with cells for 1hr at RT in a humid chamber. After incubation, slides were washed twice with fish gelatin and once with PBS before mounting with Vectashield mounting medium with DAPI. Excess mounting medium was removed and the coverslips were sealed using nail varnish.

### Microscopy

Z stack images were acquired on a Zeiss AX10 microscope equipped with AxioCam MRm charge-coupled device camera using AxioVision software (Carl Zeiss International, UK). Best exposure time for each field and channel was manually determined and kept fixed among experiments. Further image editing and refinement was achieved through Fiji/ImageJ.

### RNA extraction

Cells grown on T25 flasks (Nunc) were washed twice in PBS and lysed directly with 1ml of Trizol reagent (Life Technologies). Samples were incubated for 5 min at 25°C, cellular lysates were collected and transferred into 1.5ml RNase-free Eppendorf tubes. 0.2 volume of chloroform was added to each sample and mixed by vigorous shaking for 15 s. Samples were incubated for 2 min at 25°C and centrifuged at 12,000g for 5 min at 4 °C. The upper aqueous phase was transferred into a clean tube and an equal volume of isopropanol was added. Samples were incubated for 10 min, and RNA was pelleted at 12,000g for 10 min at 4°C. The pellets were washed once in 1ml of 75% ethanol and then air-dried for 5-10 min and resuspended in 50-100μL RNase-free water. Contaminating DNA was removed using the Ambion DNA-free DNase Treatment kit (Life Technologies) according to the manufacturer’s instructions. cDNA was generated with pd(N)6 random hexamers (GE Healthcare) using SuperScript III reverse transcriptase (Life Sciences) according to the manufacturer’s instructions.

### Genomic DNA extraction

Cells from a confluent 90-140mm petri dish were harvested and resuspended in 5-10ml lysis buffer (10 mM NaCl, 10 mM Tris-HCl pH 7.5, 10 mM EDTA-NaOH pH 8.0, 0.5% Sodium lauroyl sarcosinate) with proteinase K added to a final concentration of 100μg/ml. Samples were incubated overnight at 55°C, and then genomic DNA was phenol/chloroform extracted and purified using 15 ml MaXtract High Density Tubes (Qiagen). Genomic DNA was precipitated with 1/25 volume of 5M NaCl and 2.5 volume of ice-cold 100% ethanol. High molecular weight genomic DNA was spooled and transferred to a new Eppendorf tube containing 1ml of 70% ethanol. DNA was pelleted, air dried and resuspended in 300-400μl 10 mM Tris pH 8.5. DNA concentration was measured by Nanodrop.

### Nuclear extraction

Nuclear extracts for Western blot analysis of CRISPR/Cas9-mediated SmcHD1 mutant cell lines were prepared essentially according to a method described previously^58^. Briefly, cells were trypsinised and washed in PBS and then resuspended in 10 packed cell volumes of buffer A (10mM HEPES pH 7.9, 1.5mM MgCl2, 10mM KCl with 0.5 mM DTT, 0.5 mM PMSF, complete protease inhibitors (Sigma) added fresh, and incubated on ice for 10 mins. Cells were recovered by centrifugation at 1500g for 5 min at 4°C. Cells were then lysed in 3 volumes of buffer A+0.1% NP-40 and incubated on ice for another 10 min. Nuclei were collected by centrifugation at 400g, 5 min at 4°C and washed once in five volumes of PBS with protease inhibitors. Recovered nuclei were resuspended in one volume of buffer C (5mM HEPES pH 7.9, 26% glycerol, 1.5mM MgCl2, 0.2mM EDTA, 250mM NaCl with complete protease inhibitors and 0.5mM DTT added fresh). Salt concentration was increased to 350mM NaCl and extraction was performed on ice for 1 hour with occasional agitation. Nuclei were pelleted at 16,000g for 20 mins at 4°C and supernatants were collected as nuclear extracts. Concentration of the extracts was measured using Bradford assay (Bio-Rad) according to manufacturers’ instructions. Samples were stored at −80°C until use.

### Western blotting

Samples were diluted in 6xSMASH buffer (375mM Tris.HCl pH 6.8, 35% Glycerol, 12% SDS, 0.1% bromophenol blue, 4.3M β-mercaptoethanol), boiled for 10 min at 95°C, separated on 6% polyacrylamide gel and transferred onto a nitrocellulose membrane by wet transfer in Tris/glycine buffer (100V for 70 min at 4°C). Membranes were blocked in TBST buffer (100mM Tris-HCl pH 7.5, 0.9% NaCl, 0.1% Tween 20, 5% w/v Marvel milk powder) for 1 hour at room temperature and then incubated with primary antibodies overnight at 4°C with gentle rocking. Membranes were washed three times for 10 min with TBST and incubated for 1hr with secondary antibody conjugated to horseradish peroxidase or IRDye 800CW anti-rabbit IgG (Li-COR). After washing 3 times for 10 minutes with TBST and one 10 min wash with PBS, bands were visualised either using ECL (GE Healthcare) or on Odyssey Fc Imaging System (Li-COR).

### Bisulfite treatment and whole genome bisulfite sequencing (WGBS)

Genomic DNA and RNA were extracted from MEF cultures using AllPrep DNA/RNA/Protein Mini Kit (Qiagen) according to the manufacturer’s instructions. Concentration of DNA was measured on Nanodrop and adjusted to 20ng/µl after which the DNA was sheared to 100-500bp fragments on Covaris sonicator with the following settings: 10% duty cycle, 200 bursts/sec, intensity 4.0, 80sec duration; mode – frequency sweeping). Fragment ends were repaired, A-tailed and ligated with PE Illumina methylated adapter oligos using NEBNext DNA Library Prep Master Mix Set for Illumina (NEB). Adaptor-ligated gDNA fragments were subjected to bisulfite conversion using two-step modification procedure with Imprint™ DNA Modification Kit (Sigma). Converted and purified DNA was amplified for 9-12cycles with KAPA HiFi HS Uracil enzyme mix (Anachem). Precise concentration and size of the libraries were determined by qPCR with universal Illumina primers and also on Agilent 2100 Bioanalyzer (Agilent Technologies) using High Sensitivity DNA assay. Libraries were sequenced on HiSeq2000 system (Illumina) using 100bp Paired End protocol.

### Chromatin RNA sequencing

WT, SmcHD1 null and acute SmcHD1 null MEFs were grown on 2×145mm petri dishes in EC10 medium to semi-confluency. RNA-Seq enriching for chromatin-bound nuclear RNA was performed according to a modified chromatin RNA-Seq protocol^59^. The cells were trypsinised, collected in EC10 medium and cell number was counted on LUNAII cell counter (Logos Biosystems). 1×10^7 cells for each line were spun down, resuspended in 12ml of ice cold PBS and supplemented with 2.5×10^6 SG4 drosophila cells for calibration. Cells were spun down at 500g, 5 min at 4°C and cell pellets were resuspended in 800µl of HLBN hypotonic buffer (10 mM Tris-HCl pH 7.5, 10 mM NaCl, 2.5 mM MgCl_2_, 0.05% NP40). 480 µl of buffer HLBNS (HLBN, 25% sucrose) was carefully under-layered to create sucrose cushion, and nuclei were isolated by centrifugation for 5 min at 1000g at 4°C. Supernatant containing cytoplasmic debris was discarded and the nuclear pellet was re-suspended in 100 µl of ice-cold buffer NUN1 (20 mM Tris-HCl pH 7.9, 75 mM NaCl, 0.5 mM EDTA, 50% glycerol; 1 mM DTT and cOmplete EDTA free protease inhibitors (Sigma) added fresh). Nuclei were lysed in 1200 µl of ice-cold lysis buffer NUN2 (20 mM HEPES pH7.6, 300 mM NaCl, 7.5 mM MgCl_2_, 0.2 mM EDTA, 1 M urea, 1% NP40; 1 mM DTT) during 15min incubation on ice and RNA-bound chromatin was pelleted at 16000 g for 10min at 4°C. Chromatin-RNA pellet was re-suspended in 200 µl of high salt buffer HSB (10 mM Tris-HCl pH 7.5, 500 mM NaCl, 10 mM MgCl2). DNA and proteins were digested with Turbo DNAse (Life Sciences) and proteinase K (10 mg/ml, ThermoFisher, nuclease free), incubating on ThermoMixer at 37°C for 10 min and 30min, respectively. RNA was extracted with 1 ml of TRIzol (Life Sciences) according to the manufacturer guidelines. RNA was dissolved in 1xTURBO DNAse buffer, digested with TURBO DNAse for 30 min at 37° C on a ThermoMixer and extracted with TRIzol. RNA was washed three times with 75% ethanol, dissolved in water and quantified using a NanoDrop (ND-1000).

RNA quality was checked with RNA 6000 Pico Chip (Agilent Technologies) on Agilent 2100 Bioanalyser (Agilent Technologies). Samples were depleted of ribosomal RNA with Ribo-Zero Gold Kit (MRZG12324, Illumina) according to manufacturer’s guidelines. RNA was isolated from 3 biological replicates. RNA-Seq libraries were prepared with NEBNext^®^ Ultra™ Directional RNA Library Prep Kit for Illumina (E7420S) using NEBNext^®^ Multiplex Oligos for Illumina for multiplexing (E7335S and E7500S). Libraries were sequenced on HiSeq2000 and NextSeq500 using NextSeq 500 High-Output Kit: 1 lane, 150 cycles, 75 bp paired end sequencing (Illumina).

### Chromatin immunoprecipitation sequencing

WT, SmcHD1 null acute SmcHD1 null MEFs were grown on 5-8×145mm petri dishes in EC10 medium to semi-confluency. Cells were trypsinised, washed in PBS and counted. For SmcHD1 and Rad21, 5×10^7^ cells were then cross-linked in 2µM Ethylene glycol-bis(succinic acid N-hydroxysuccinimide ester) (EGS, Sigma) in PBS at RT for 1hr followed by 15 min cross-linking in 1% formaldehyde. For H3K27me3 and CTCF 1×10^7^ cells were crosslinked in 1% formaldehyde alone for 15 min at RT. Formaldehyde was quenched by addition of glycine to a final concentration of 125mM, and incubation at RT for 3min. Cells were spun down at 700rpm for 4 min at 4°C and lysed in 50mM HEPES pH 7.9, 140mM NaCl, 1mM EDTA, 10% glycerol, 0.25% Triton X100 and 2% NP40. Cellular lysis was assisted by 20 strokes of a large clearance pestle in Dounce grinder, with subsequent incubation for 10 min at 4°C with constant rotation. Released nuclei were spun down and washed once in 10mM Tris-HCl pH8.0, 200mM NaCl, 1mM EDTA and 0.5mM EGTA. Nuclei were lysed by addition of 0.1% sodium deoxycholate and 0.5% N-lauroylsarcosine. All solutions contained cOmplete EDTA-free protease inhibitors (Sigma) added fresh. Chromatin was sonicated on BioRuptor sonicator (Diagenode) to produce fragments of approximately 500 bp. Triton X100 was added to a final concentration of 1%, and insoluble pellet was removed by centrifugation at maximum speed for 10 min at 4°C. Chromatin was aliquoted and stored at −80C until use.

Immunoprecipitation was performed overnight at 4°C with 3-5µg of specific antibody and 100 µl of chromatin corresponding to 1×10^5^ (H3K27me3, CTCF) or 5×10^5^ cells (Rad21, SmcHD1) in 20 mM Tris-HCl pH 8.0, 1mM EDTA, 150 mM NaCl and 1% triton-x-100, with proteinase inhibitors. Antibody-bound chromatin was isolated on rProtein A Sepharose Fast Flow beads (GE Healthcare) that had been blocked for 1 h at 4°C with 1 mg/ml bovine serum albumin (NEB) and 1 mg/ml yeast tRNA (Sigma). Agarose beads with immunoprecipitated material were washed thoroughly in low salt buffer (LSB, 20mM Tri-HCl pH8.0, 2mM EDTA, 150mM NaCl, 0.1% SDS, 1% Triton X100), high salt buffer (HSB, the same as LSB but with 500mM NaCl), LiCl buffer (10MM Tris-HCl pH8.0, 1mM EDTA, 0.25 M LiCl, 1% NP40, 1% deoxycholate) and twice in TE buffer, all with proteinase inhibitors. Immunoprecipitated material was eluted from beads in elution buffer (0.1M NaHCO_3_, 1% SDS) with shaking on Thermomixer at RT for 30min. Beads were removed by centrifugation, eluted chromatin was reverse cross-linked and treated with RNAse and proteinase K in the presence of 200mM of NaCl at 37°C for 2 hrs followed by 65°C overnight with shaking at 800rpm for 1 min every 2 min. DNA was purified using ChIP DNA Clean and Concentrator kit (Zymo Research) according to the manufacturer’s instructions.

DNA size was assessed on Bioanalyzer (Agilent) and DNA was post-sonicated on Bioruptor Pico sonicator for 18-20 cycles of 30s on/30s off if necessary. DNA Concentration was quantified using PicoGreen^®^ dsDNA Quantitation Kit (Molecular Probes), Bioanalyzer High sensitivity DNA assay and/or Qubit dsDNA HS assay kit.

NEBNext (H3K27me3) or NEBNext Ultra II (CTCF, Rad21, SmcHD1) DNA Library Prep Kits for Illumina were used to prepare libraries for sequencing, following the manufacturer’s instructions. End-repaired, A-tailed and adapter-ligated libraries were amplified for 7-11 cycles depending on the initial DNA amount. Indexed libraries were quantified, normalised and pooled for sequencing either on Illumina HiSeq2000 50bp paired end run (H3K27me3, 8 samples on 3 lanes) or on Illumina NextSeq 550 System (CTCF, Rad21, SmcHD1, 8-12 libraries per flowcell).

### Repli-seq

Repli-seq was performed as described^60^. Briefly, asynchronously proliferating cells were flow-sorted based on their DNA content (Hoechst 33342) into G1 and S phase fractions (Fig.S6) into ice-cold PBS. Genomic DNA was isolated immediately after flow-sorting with Quick-gDNA™ MiniPrep (Zymo Research). Sequencing libraries were prepared with Illumina TruSeq DNA PCR-Free whole genome sequencing kit and sequenced on NextSeq 550 with 150 cycles High Output kit using paired end protocol.

### HiC

Hi-C was performed as described^61^. For each Hi-C library 25 × 10^6 cells were cultured in EC10 media. The day of harvesting, cells were incubated in 22.5 mL of fresh media without serum, and cross-linked by adding 625 μL of 37% formaldehyde (1% final concentration). Plates were immediately mixed thoroughly after formaldehyde addition, and subsequently rocked every 2 minutes for exactly 10 minutes at RT. The cross-linking was quenched by addition of 1.25 mL of 2.5 M glycine. After 5 minutes at RT, plates were placed on ice for an additional 15 minutes. Cells were finally harvested by a cell lifter, transferred to a 15 mL falcon tubes, and centrifuged at 800 g for 10 min (4°C). The cell pellet was snap-frozen in liquid nitrogen, and stored at −80°C until use. Hi-C libraries were then sequenced on a HiSeq4000 (paired end reads of 100 bases each). Two biological replicates of WT and SmcHD1 null MEFs were sequenced in separate lanes, each yielding ∼350 million reads per replicate.

## BIOINFORMATIC ANALYSIS

### Allele-specific alignment

All NGS data were obtained from interspecific FVB- CAST/EiJ cells which enabled allele-specific analysis. Assigning of reads into one of the parental genomes was performed in two stages.

First reads were mapped to mm10 reference genome with the aligner optimal for the assay. To minimise mapping biases due to differences in the similarity of FVB/Cast genomes to the reference genome all known SNP loci were N-masked before generating appropriate genome index files. SNP coordinates were obtained from The Sanger Institute (ftp://ftp-mouse.sanger.ac.uk/REL-1505-SNPs_Indels/).

Second using SNPsplit programme^62^ and confident SNPs extracted from the file mentioned above reads were sorted into paternal, maternal and unassigned subsets. The procedure in our experimental set up allowed us to assign ∼30% reads to parental genomes.

### Chromatin RNA-seq

Reads were mapped with STAR 2.5b aligner^63^ with index generated from FVB-Cast SNP N-masked genome. Counts per gene were obtained with Htseq-count^64^.

Initial analysis showed that there is high correlation between chromosome copy number and gene expression values on distinct parental chromosomes therefore HTSeq counts were normalised for chromosomal copy number differences estimated from comparison of ChIPseq input values and repli-seq alignments.

Differential expression analysis was performed with DESeq2^65^, a package from the R Bioconductor project. Results were further processed, analysed and visualized with custom R scripts using commonly used base and Bioconductor packages.

### ChIP-seq

ChIP-seq reads were mapped with bowtie2 with SNP N-masked genome index and sorted with SNP-split to separate subsets of reads originating from parental genomes. For SmcHD1, CTCF and Rad21 ChIP-seqs, macs2 was used to detect significant enrichments (narrow peaks, q-value < 0.05). Peak calling was performed on unsorted reads. Allele-specificity of distinct peaks was determined by calculating the ratio of allele-specific reads overlapping the peak in maternal and paternal genome. Peak were considered exclusively specific to either paternal or maternal chromosome if they contained more than 90% of all input corrected reads.

Chromosome-wide SmcHD1 and H3K27me3 ChIP-seq data were obtained by averaging log2(IP/input) values calculated for 500 kb bins within 10 kb intervals.

H3K27me3 depletion domains were defined as regions with intervals of mean log2(IP/input) <0, adjacent regions with negative values separated by no more than 20 kb were merged.

### Hi-C

Hi-C reads were mapped with HiCUP pipeline^66^ using bowtie2 indexes based on the FVB-Cast SNP N-masked mm10 genome. Valid sets of Hi-C ditags were obtained after removing uninformative reads (re-ligations, dangling ends, etc), as well as exact duplicates. Hi-C ditags were then sorted based on confident SNP content with SNPsplit into following groups: FVB-FVB, FVB-unassigned, Cast-Cast, Cast-unassigned, FVB-Cast, unassigned-unassigned. For allele-specific analysis subsets “FVB-FVB” and “FVB-unassigned” were merged into one “FVB” and similarly subsets “Cast-Cast” and “Cast-unassigned” were merged into subset “Cast”. For comparative analysis of the WT and mutant cells, Hi-C data interaction number were down-sampled to the size of the smallest sample and further equalized per chromosome to normalize for differences in DNA copy number. Interactions were further binned into 1 Mb intervals genome-wide and 100kb and 50 kb bins chromosome-wide with juicer tools^67^. Matrices with interactions for distinct chromosomes were balanced separately with the Knight-Ruiz balancing algorithm implemented in juicer tools, that ensures that each row and column of the contact matrix sums to the same value. TAD borders were called with TADtool^68^. Insulation scores were calculated withthe <u>matrix2insulation.pl</u> script developed in Dekker Lab based on 50 kb balanced matrices with the following options: -is 500000 -ids 200000 -im imean). The <u>matrix2insulation.pl</u> script is available from https://github.com/dekkerlab/cworld-dekker.

Heatmaps and sums of interaction numbers for specific parts of chromosomes were generated based on balanced Hi-C matrices with HiTC Bioconductor package^69^.

### Whole Genome Bisulfite sequencing

Whole Genome Bisulfite libraries were mapped with Bismark^70^, and deduplicated. Further reads were allele-specifically sorted and processed with bismark_methylation_extractor to retrieve methylated cytosines. Cytosines covered by at least 3 reads in allele-specific alignment were used for further analysis.

### Repliseq

Repli-seq reads were mapped with bowtie2. Obtained alignments were sorted into parental genome specific subsets and subsequent analysis were performed as described^60^. In brief, reads from S and G1 cell fractions were analysed in 100 kb and 500 kb bins and for each bin read number normalized S/G1 ratio was plotted. Z-scores of S/G1 ratios were further used for detailed analysis and visualization of chromosome-wide replication timing profiles.

### Oligonucleotides used in this study

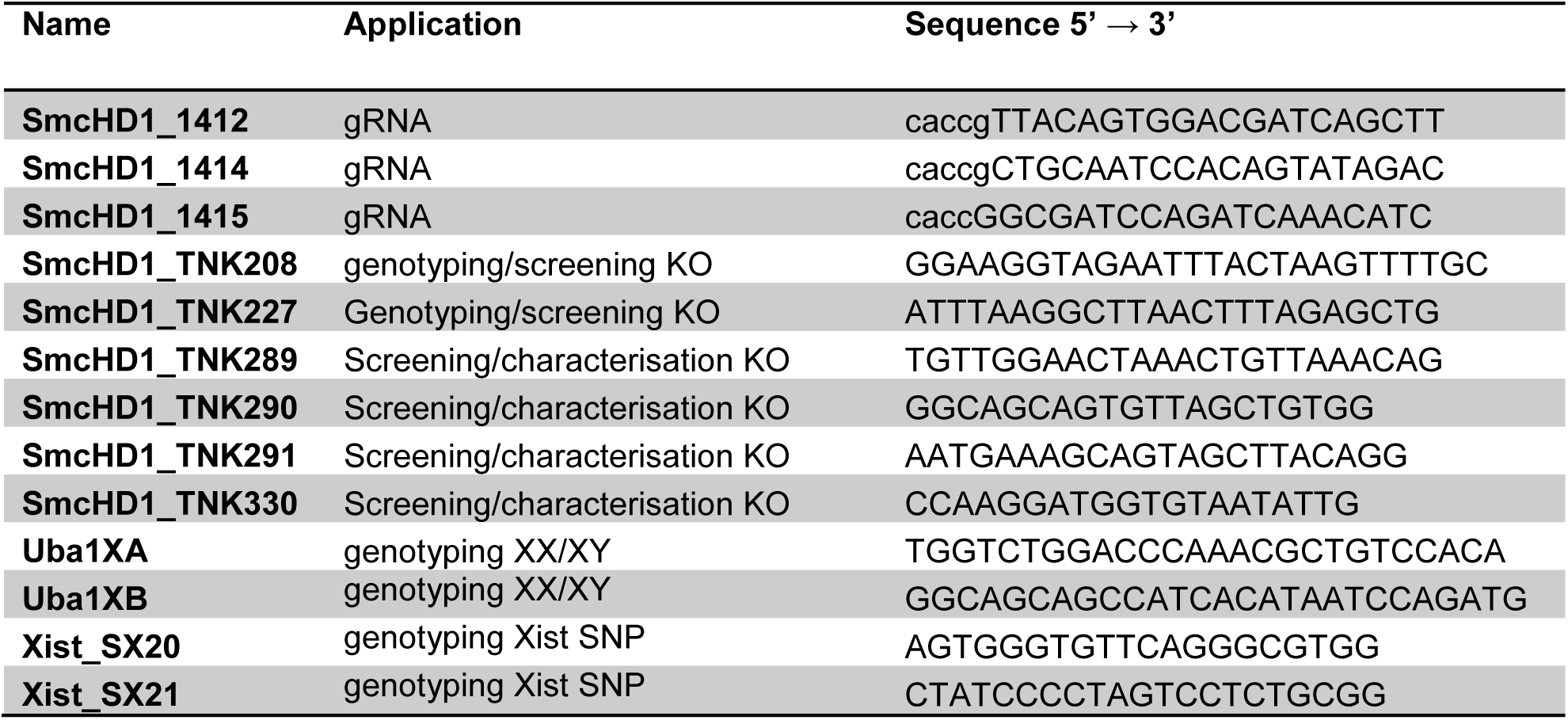

### Antibodies used in this study

| Antibody | Raised in | Source | Reference/<br>Cat. No. | Dilution for<br>IF | Dilution<br>for WB | Used per<br>IP |
| --- | --- | --- | --- | --- | --- | --- |
| **H3K27me3** | Rabbit | Diagenode | C15410069 | - | - | 5µg |
| **IgG** | Rabbit | Sigma | M7023 | - | - | 5µg |
| **SmcHD1 anti-**<br>**mouse** | Rabbit | Raised in<br>house | Brideau et<br>al, 2015 | 1:500 | 1:1,000 | 10µl |
| **Rad21** | Rabbit | Abcam | ab992 | - | - | 5µg |
| **CTCF** | Rabbit | Millipore | 07-729 | - | - | 5µl |
| **H2AK119u1** | Rabbit | Cell<br>Signalling | mAb #8240 | 1:1,000 | - | - |
| **H3K27me3** | Mouse | Active Motif | 61017 | 1:1,000 | - | - |
| **Alexa 568 anti-**<br>**mouse IgG** | Goat | Life<br>Technologies | A11031 | 1:500 | - | - |
| **Alexa 488 anti-**<br>**rabbit IgG** | Goat | Life<br>Technologies | A11008 | 1:500 | - | - |
| **Anti-HDAC3 [Y415]** | Rabbit | Abcam | mAb<br>ab32369 | - | 1:5,000 | - |
| **Anti-rabbit IgG,**<br>**HPR** | Donkey | Amersham | NA934 | - | 1:2,000 | - |
| **IRDye 800CW anti-**<br>**rabbit IgG** | Goat | Li-COR | 926-32211 | - | 1:15,00 |  |

## DATA AND SOFTWARE AVAILABILITY

The accession number for the data (…….-seq) reported in this study were deposited in GEO under GSEXXXXX.

## CONTACT FOR REAGENT AND RESOURCE SHARING

Further information and requests for resources and reagents should be directed to the corresponding author, Prof. Neil Brockdorff

---

**Figure S1.**
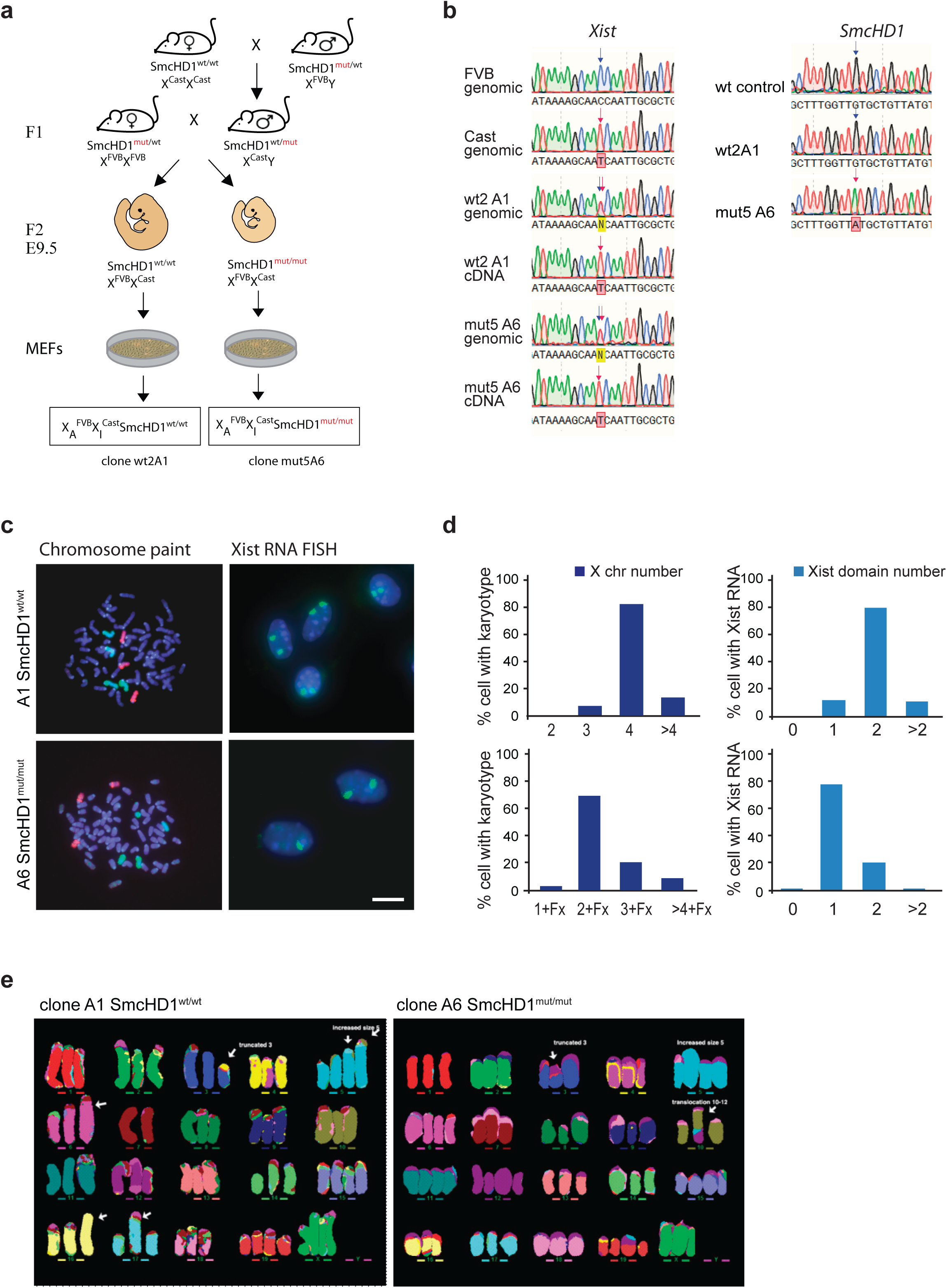
A MEF cell model for allele specific analysis of Xi/Xa. a. Schematic indicating genotypes and direction of crosses used to generate F2 embryos for MEF lines. Established MEF lines were genotyped and subcloned to isolate clonal lines carrying either castaneus (cast) or domesticus (FVB) Xi. b. Left; sequencing traces of FVB, castaneus and F2 progeny Xist PCR fragments encompassing SNP region. Genomic and cDNA traces are shown to identify the origin of Xi for the cell lines studied. SNP is indicated in blue (C, FVB) or red (T, Cast) arrow. Right; sequencing traces of WT and mutant SmcHD1 PCR fragments. WT (G, blue arrow) and SmcHD1 null (A, red arrow) SNPs are indicated. c. Karyotype analysis of the WT (clone A1, top) and SmcHD1 null (clone A6, bottom) cell lines. Total number of X chromosomes detected by Chr X paint is shown in green and Chr 8 in red. Xist RNA FISH shows the number of inactive X chromosomes. Bar, 10 µm d. Quantification of the karyotype and Xist RNA FISH analyses. At least 32 metaphase spreads and over 130 cells were scored for karyotyping and RNA FISH, respectively. e. Examples of multiplex in situ hybridization (M-FISH) karyotypes of the WT (left) and SmcHD1 null (right) cell lines.

**Figure S2.**
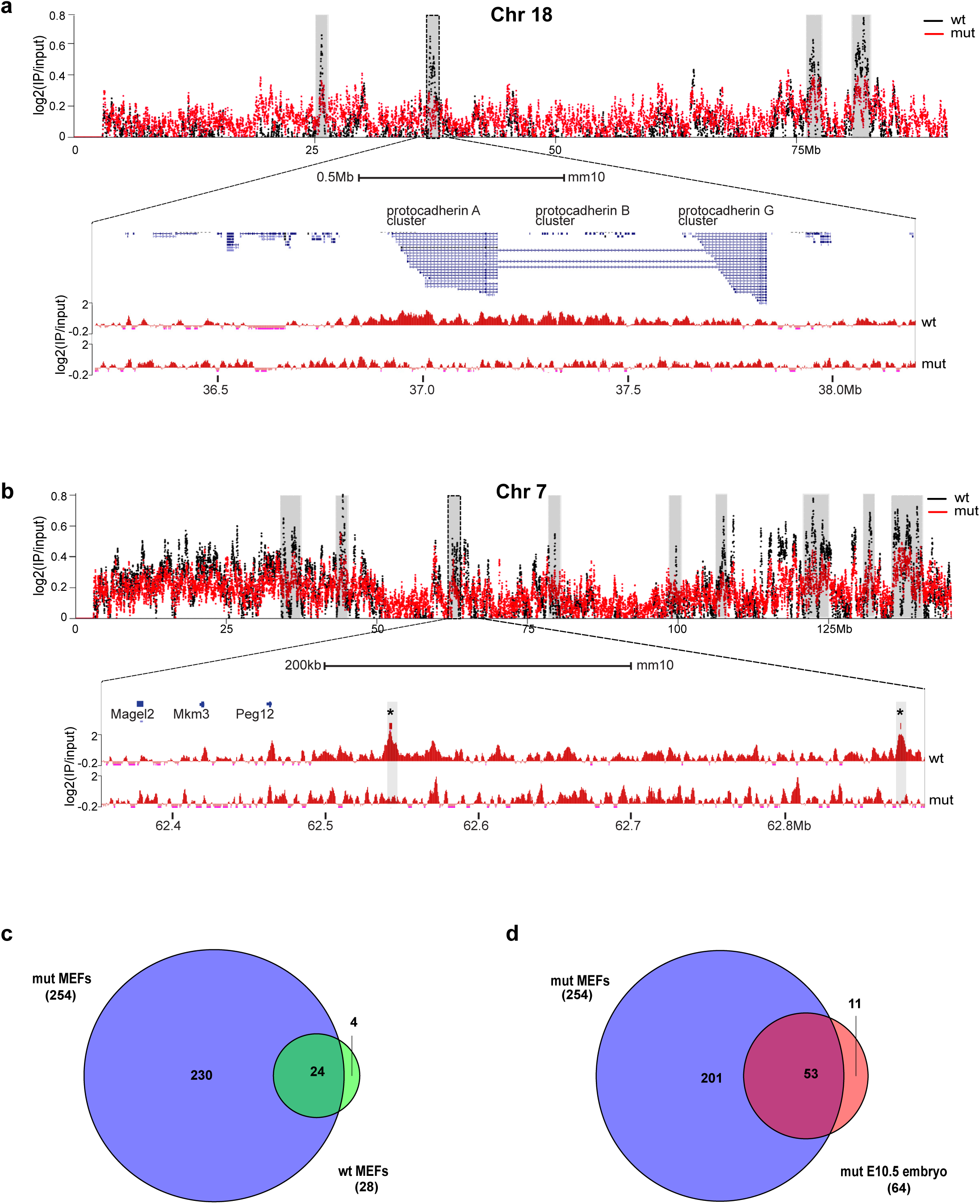
Analysis of SmcHD1 occupancy and transcription on Xi. a. Chromosome-wide profiles of SmcHD1 occupancy on Chr 18 depicted as allele-specific ChIP-seq log2 ratios of IP to input per 500 kb binned into 10 kb intervals. Profiles from WT (black) and SmcHD1 null MEFs (red) are presented. Regions enriched for SmcHD1 are shaded with magnification shown for the protocadhedrin cluster. b. Chromosome-wide profiles of SmcHD1 on Chr 7 presented as in (a). Magnified region covers the imprinted gene cluster Magel2, Mkrn3, and Peg12. Specific peaks indicated with asterisks were also reported in an independent stud y^37^. c. Genes expressed from Xi in SmcHD1 null (mut) and WT MEFs. Total number for each cell type is shown in brackets. d. Comparison of genes expressed from Xi in SmcHD1 null MEFs in this study, and in embryonic E10.5 cells with the same mutation, as published previously^34^. Total number for each study is shown in brackets.

**Figure S3.**
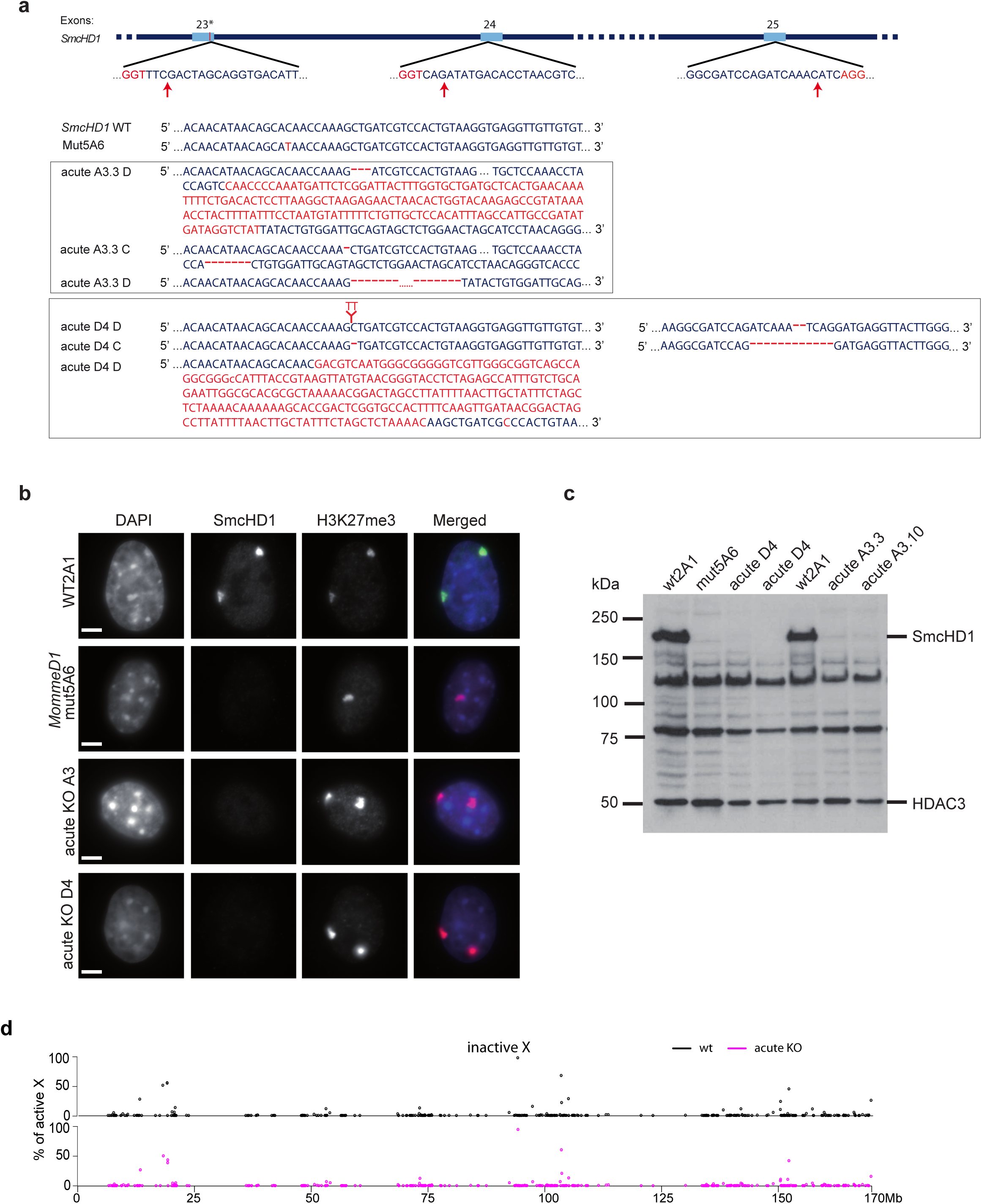
Generation and characterisation of acute SmcHD1 null MEF lines. a. Strategy for acute CRISPR-Cas9-mediated mutagenesis of SmcHD1 gene. Relative position of the MommeD1 point mutation is indicated by the vertical red bar at the distal part of the exon 23. The positions of guide RNAs relative to SmcHD1 exons are shown, with PAM sequences (red font) and predicted cleavage sites (red arrows) indicated. Two sgRNAs for exons 23 and 24 were used to generate a mutant clone A3.3 and two sgRNAs for exons 23 and 25 were used to generate clone D4. An alignment of WT with mutated sequences of domesticus (D) and castaneus (C) alleles of the clones A3.3 and D4 is shown below. As both clones are triploids, three independently mutated alleles are shown for each clone. Deletions are shown by red dashes and insertions/mutations are shown in red font. b. Immunofluorescence analysis of SmcHD1 and H3K27me3 in WT, SmcHD1 null and acute SmcHD1 null MEF lines. Images show representative examples of cells with H3K27me3 Xi domains in all cell lines and loss of SmcHD1 Xi domains in SmcHD1 null and acute SmcHD1 null mutant cells. Scale bar is 5 μm c. Western blot analysis of acute SmcHD1 null MEF lines in comparison with WT and SmcHD1 null MEFs. HDAC3 is included as a loading control. Bands of approx. 80kDa and 130kDa are non-specific proteins recognised by the SmcHD1 antibody. Molecular weight markers are shown on the left. d. Gene de-repression on WT and acute SmcHD1 null Xi, presented as % of Xa expression.

**Figure S4.**
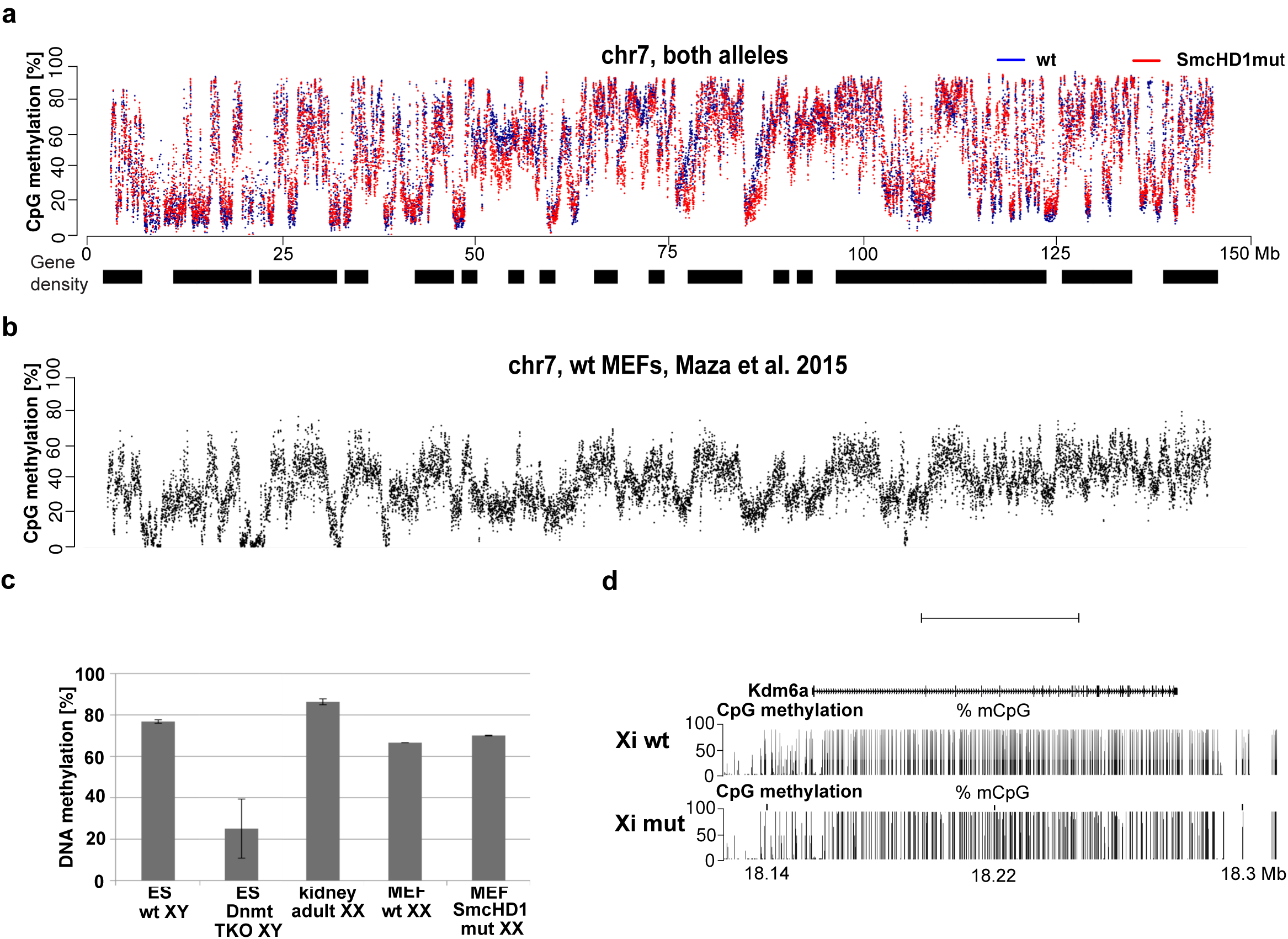
DNA methylation in WT and SmcHD1 null MEFs. a. DNA methylation profile of Chr 7 in WT and SmcHD1 null MEFs plotted as % of mCpG averaged within 10 kb bins. b. DNA methylation profile of Chr 7 using published MEF WGBS data ^38^. c. Global DNA methylation level of MEF lines used in this study and embryonic stem (ES) cells or adult tissue determined by HPLC. TKO denotes Dnmt1, Dnmt3a, Dnmt3b triple knockout ES cells. d. Example of DNA methylation profile of known escapee, Kdm6a (UCSC screen shot).

**Figure S5.**
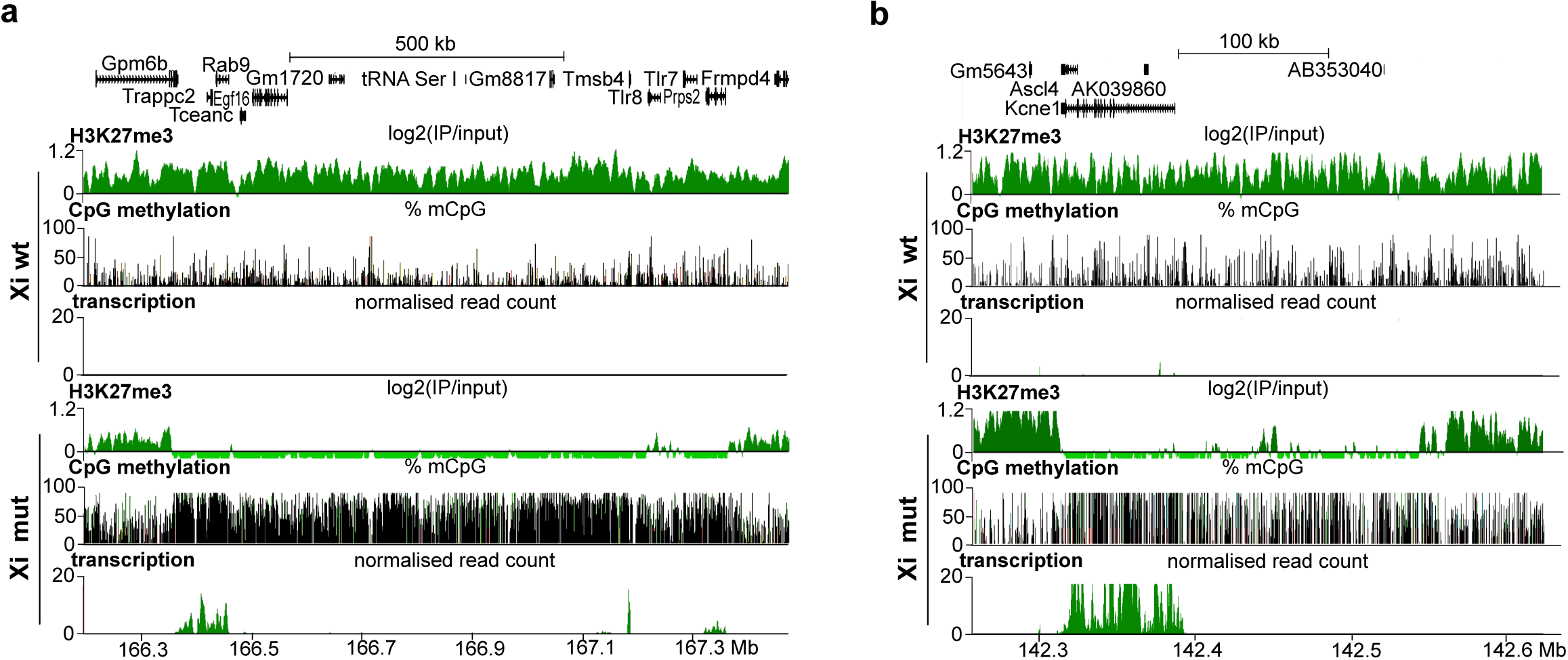
Domains depleted of H3K27me3 in SmcHD1 null MEFs. a. Example showing large domain that is H3K27me3 depleted on the SmcHD1 null Xi and that encompasses both transcribed and silent genes. b. An example of an H3K27me3 depleted domain in which in which there are no annotated genes or detectable transcription.

**Figure S6.**
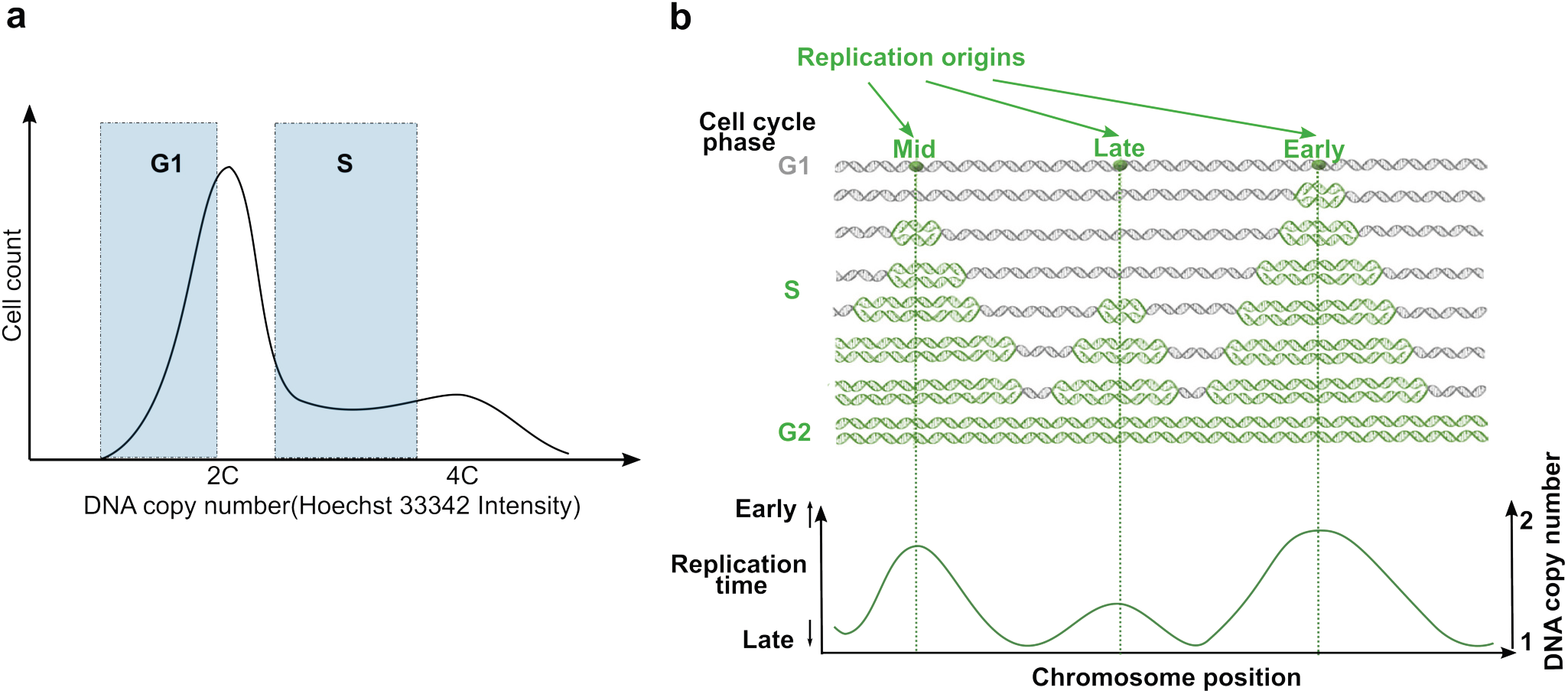
Generation of genome-wide replication timing profiles using Repli-seq. a, Asynchronously proliferating cells are flow-sorted based on their DNA content (Hoechst 33342) into G1 and S phase fractions. b, All cells in S-phase have early replicating regions of DNA copied, but regions replicating later are doubled only in cells which are more advanced in genome replication. Therefore in DNA extracts from asynchronous cell populations, early replicating DNA is most abundant. The later replication occurs, the lower is the copy number of the DNA sequence. Thus, S-phase DNA copy number profile obtained with high-throughput sequencing (coverage profiles, PCR-free library prep), normalised with an analogous profile for G1 phase from the same cell population, allows the genome-wide replication timing profile to be obtained.

**Figure S7.**
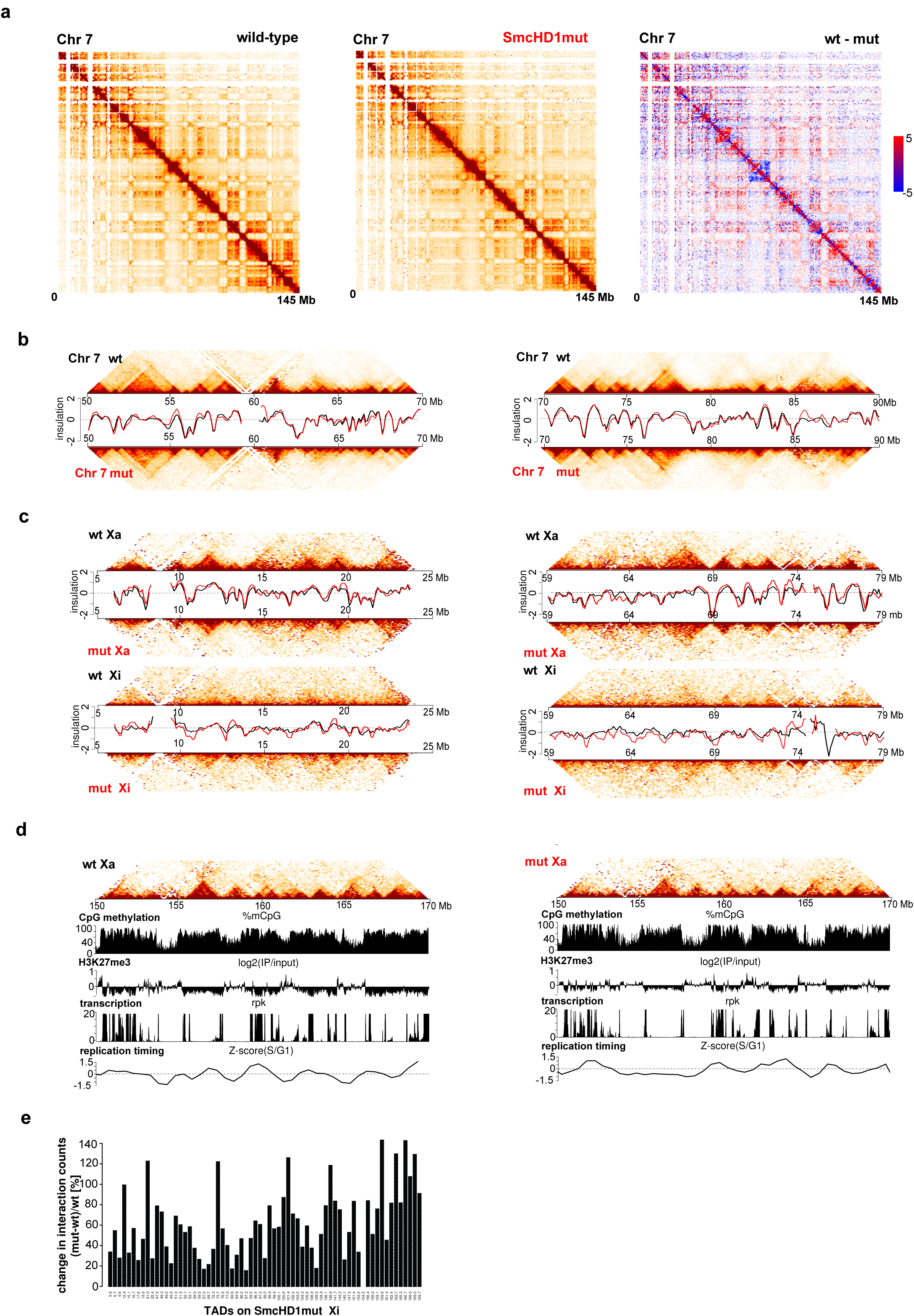
Higher-order structure of autosomes and Chr X in SmcHD1 null and WT MEFs. a. Heatmap depicting Hi-C interactions for Chr 7 of WT and SmcHD1 null MEFs. Interaction matrices were generated from Hi-C interactions normalised for the available read number per chromosome, ICE-balanced and binned into 500 kb bins. Right side of the panel presents difference between normalized interaction counts of WT and SmcHD1 null MEFs. b. Heatmaps presenting local chromatin conformation of two representative 20 Mb regions on Chr 7. Interactions binned into 50 kb bins. c. Local Hi-C interactions within two additional representative 20 Mb regions on Chr X. Allele-specific Hi-C heatmaps for Xa and Xi of WT and SmcHD1 null MEFs were generated as in Fig. 6D. d. Similar to Fig.6E, allele-specific Hi-C heatmaps of Xa of the WT and SmcHD1 null MEFs aligned with respective DNA methylation, H3K27me3, transcription and replication timing profiles for the distal 20 Mb of Chr X. e. Similar to Fig.6F, comparison of the interaction counts from distinct TADs on the Xi of SmcHD1 null MEFs, with the interaction counts within the same intervals on the WT Xi. The values were calculated as the difference between the counts in SmcHD1 null and WT MEFs relative to the count number in WT MEFs and presented as %. All interactions falling within TAD borders were counted.

**Figure S8.**
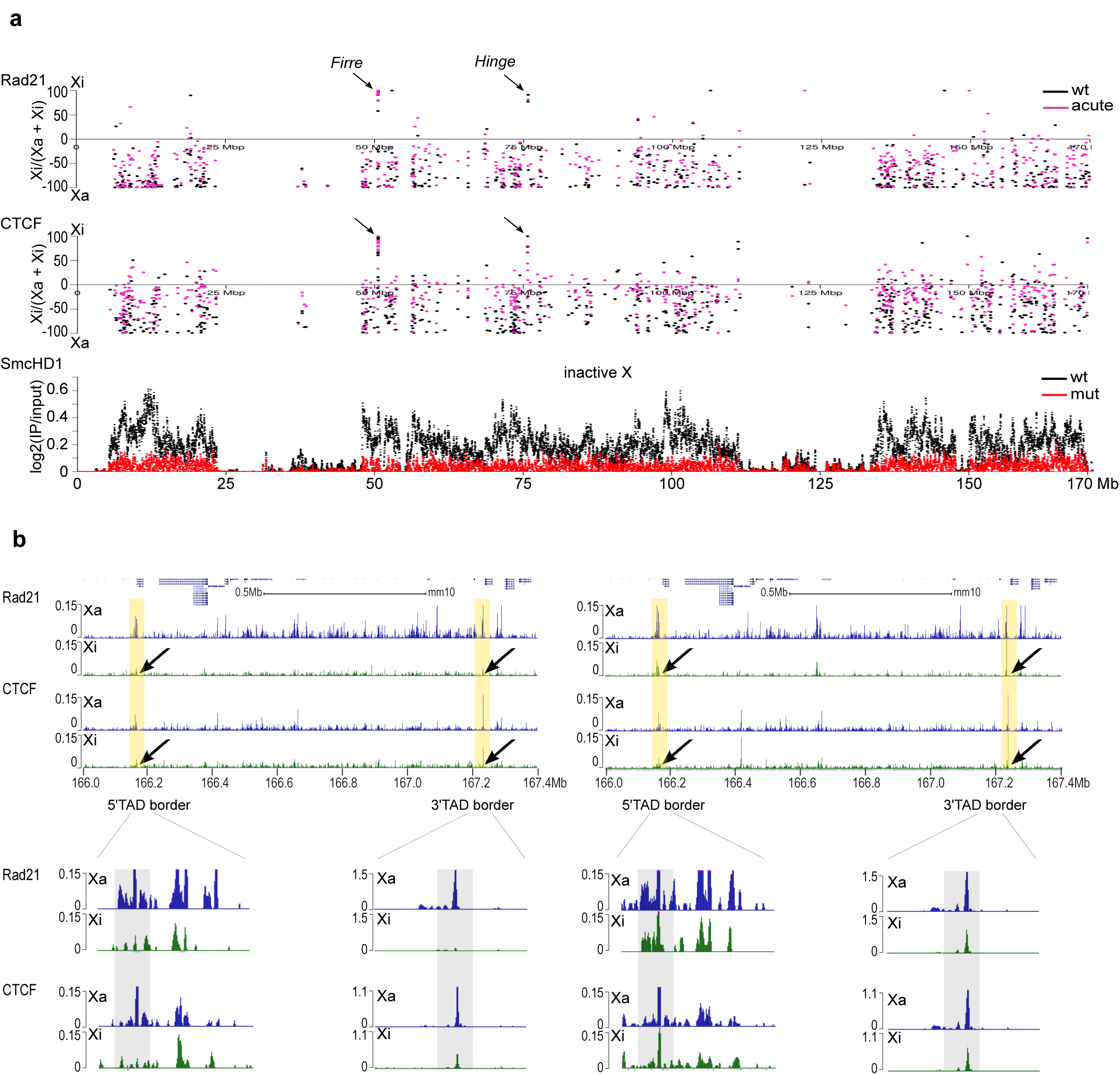
CTCF and Rad21 ChIP-seq analysis. a. Chromosome-wide occupancy of Rad21 (top panel) and CTCF (middle panel) for Xi or Xa, estimated for distinct peaks occupied in all three cell lines. Values for WT (black) and acute SmcHD1 null MEFs (red) calculated as in Fig.6a. Black arrows depict Firre and Hinge loci. Bottom track shows for reference SmcHD1 occupancy profile on the WT Xi as log2(IP/input) in 10 kb bins. b. CTCF and Rad21 distribution for the acute SmcHD1 null MEFs over the locus presented in Fig.6c. Black arrows depict CTCF/Rad21 occupancy changes within the borders of the reestablished TAD. Yellow shading highlights regions which differ in CTCF/Rad21 binding between WT and acute SmcHD1 null cells, which correlate well with external/ internal borders of the re-established TAD. Lower panels show magnified 5’ and 3’ TAD borders. Grey shading highlights TAD borders where CTCF and Rad21 peaks reappear in acute SmcHD1 null MEFs.

